# The trilobite nervous system and the early evolution of the mandibulate central complex

**DOI:** 10.64898/2026.10.05.756825

**Authors:** Sarah R. Losso, Bruno Becker-Kerber, James C. Weaver, Francesc Pérez-Peris, Javier Ortega-Hernández

**Affiliations:** Museum of Comparative Zoology and Department of Organismic and Evolutionary Biology, Harvard University, Cambridge, MA 02138, USA; Wyss Institute for Biologically Inspired Engineering, Harvard University, Cambridge, MA 02138, USA; Department of Earth and Environmental Sciences, University of Iowa, Iowa City, Iowa 52242, USA

## Abstract

Fossil central nervous systems (CNS) have illuminated several facets of total-group euarthropod evolution, including their phylogenetic affinities, the origin of the head, and their ancestral neuroanatomical organization. However, the CNS remains unknown in trilobites, the most diverse and earliest group of Palaeozoic euarthropods, but can potentially clarify fundamental questions that remain disputed despite thousands of described species and countless studies spanning over two centuries, such as their affinities relative to modern representatives. Here, we describe the CNS of the cheirurid *Anacheirurus adserai* from the Lower Ordovician Fezouata Shale. *Anacheirurus* features a brain with well-developed proto, deuto- and tritocerebral neuromeres, an anteriomedian protocerebral protrusion, a central complex, an esophageal foramen, and a ganglionated ventral nerve cord. The CNS is replicated in framboidal iron that differs in texture and composition from the exoskeleton and rock matrix. The overall organization of *Anacheirurus’* CNS most closely resembles that of Cambrian stem-group mandibulates, and critically shares the presence of a central complex with a protocerebral bridge and lateral accessory lobes with extant pancrustaceans. Our findings provide new morphological support for the hypothesis that Trilobita belongs to the mandibulate stem lineage and bring new insights into the ancestral morphology and evolutionary history of the euarthropod central complex. More broadly, our findings offer direct anatomical evidence of the key sensorimotor system innovations that originated during the Cambrian Information Revolution.

## Introduction

Trilobites are the most iconic group of extinct euarthropods due to their extensive fossil record encompassing over 22,000 described species and evolutionary history spanning over 270 million years^1^. The decay-resistant biomineralized dorsal exoskeleton of trilobites^2^, coupled with their elaborate morphologies^3,4^, versatile autecology^5–7^, well-documented developmental patterns^8,9^, and widespread paleobiogeographic distribution makes them ideal for studying early animal life near the start of the Paleozoic Era, particularly in the context of their sudden worldwide appearance during the Cambrian Explosion^10,11^. Trilobites have been the focus of extensive research, and their commonality in sites with exceptional fossil preservation has provided critical insights into the evolution of their external (e.g. appendages^5,12,13^) and internal anatomy (e.g. guts^6,14^). Despite this inordinate fondness for trilobites, their precise phylogenetic affinities relative to extant euarthropods remain as contested today as a century ago^1,15–19^, specifically whether they are more closely related to Chelicerata (e.g. horseshoe crabs, arachnids), to Mandibulata (e.g. myriapods, pancrustaceans), or occupy an earlier branching position in stem-group Euarthropoda. This complication stems from the fact that trilobite exoskeletal and appendicular morphology shares several ancestral traits with both chelicerates (e.g. lamellate gills^13^, gnathobasic protopodites^7,20^, mating claspers^5^) and mandibulates (e.g. uniramous antennae^15,16^, compound eye ultrastructure^21–23^), creating character conflict that offers phylogenetic support for either hypothesis depending on the analytical parameters^24–26^. Some of the recent morphological arguments for trilobites as stem-group euarthropods have largely relied on various interpretations of the homology of cephalic sclerites, particularly the anterior sclerite and hypostome, relative to other groups (e.g.^27–29)^. Ultimately, resolving trilobite affinities is key for reconstructing early euarthropod evolution because they represent the stratigraphically oldest representatives of this phylum^10,11^, are among the earliest complex bilaterian bioturbators in the fossil record^30^, and their phylogenetic position directly impacts that of numerous non-biomineralized antennae-bearing close relatives known from exceptional Lower Paleozoic sites worldwide^31,32^. Although numerous studies in the last decade have demonstrated that paleoneuroanatomical data contains critical phylogenetic information that can inform the affinities of problematic Cambrian total-group euarthropods^33–45^, no traces of nervous tissues have been found in trilobites to date. Here, we describe the first record of a fossilized central nervous system (CNS) in Trilobita and explore its implications for the group’s affinities and early euarthropod paleoneuranatomical macroevolution.

## RESULTS

### *Anacheirurus* internal anatomy

The studied material consists of seven specimens of the cheirurid (Phacopida) *Anacheirurus adserai* from the Lower Ordovician Fezouata Shale biota of Morocco^46^ (Fig. 1; Supplemental Table 1), a site with exceptional fossil preservation that combines faunal elements from Cambrian Burgess Shale-type biotas alongside Ordovician taxa^47^. Specimens are identified as *Anacheirurus adserai* based on their exoskeletal morphology, including: semicircular cranidium with a nearly horizontal anterior margin, a sub-rectangular glabella with sub-parallel lateral margins and rounded antero- and postero-lateral corners, three well expressed postero-medially directed glabellar furrows (S1 furrow posteriorly deflected medially), and long thoracic pleurae with a deep furrow that runs obliquely^46^. Three of the studied specimens (MCZ:IP:202504, MCZ:IP:202507, and MCZ:IP:202508) preserve remains of the digestive system (Figs. 2g,h. 3e).

**Figure 1.**
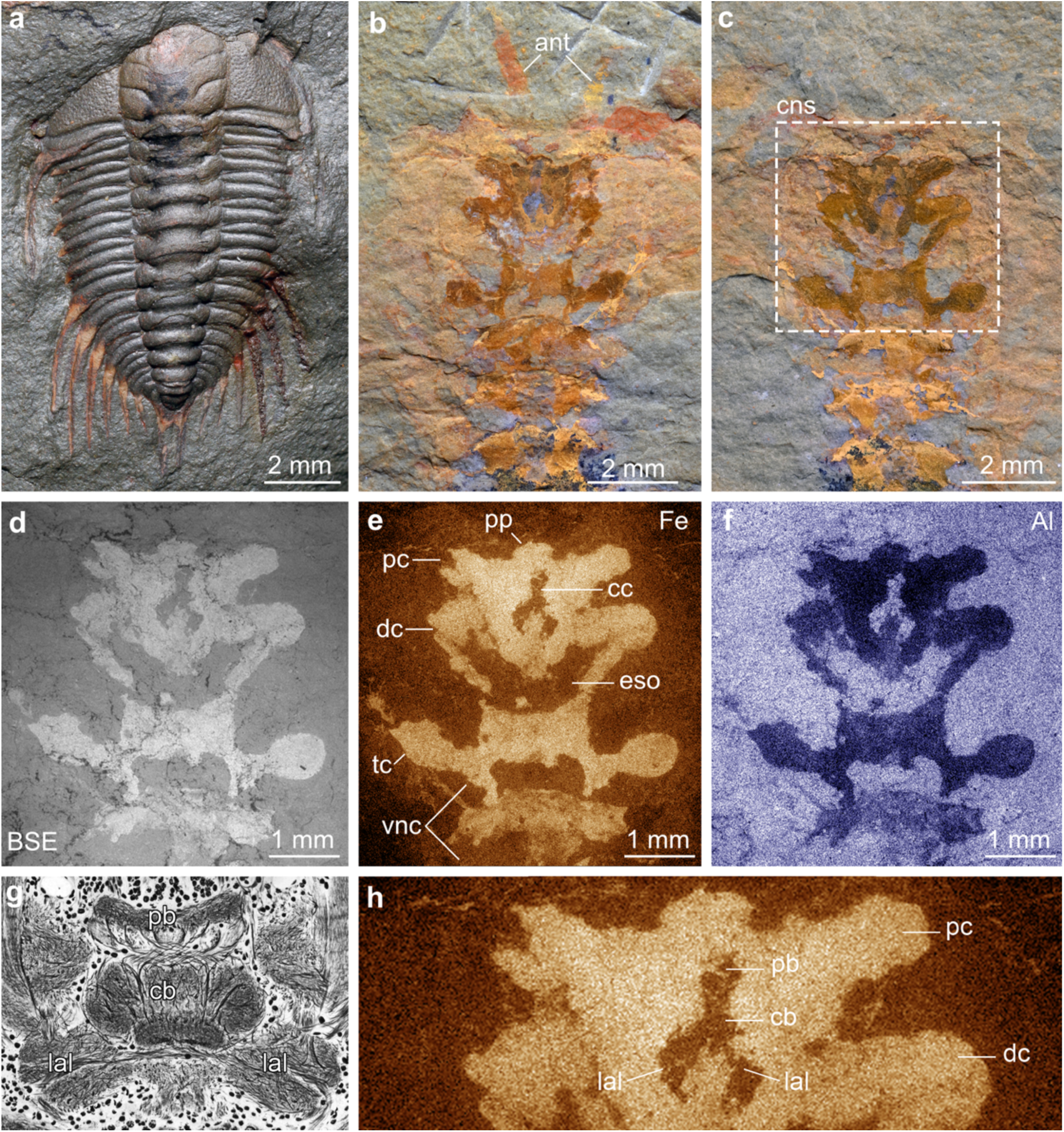
Central nervous system preservation in the cheirurid trilobite *Anacheirurus adserai* from the Lower Ordovician Fezouata Shale. **a,** YPM 530933, complete holaspid specimen showing dorsal exoskeletal morphology. **b,** MCZ:IP:202503a, specimen showing fossilized central nervous system (see fig. S1). **c,** MCZ:IP:202503b, counterpart. **d,** MCZ:IP:202503b, backscattered electron micrograph of nervous system. **e,** MCZ:IP:202503b, elemental map showing nervous system enrichment in iron. **f**, MCZ:IP:202503b, elemental map showing nervous system depletion in aluminum. **g,** Central complex of locust *Schistocerca gregaria*^54^. **h,** Magnification of MCZ:IP:202503b showing central complex visible through elemental map of iron. Institutional abbreviations: Museum of Comparative Zoology, Harvard University (MCZ); Yale Peabody Museum (YPM). Abbreviations: ant, antenna; cb, central body; cc, central complex; cns, central nervous system; dc, deutocerebrum; eso, esophageal foramen; lal, lateral accessory lobes; pb, protocerebral bridge; pc, protocerebrum; pp, protocerebral protrusion; tc, tritocerebrum; vnc, ventral nerve cord.

**Figure 2.**
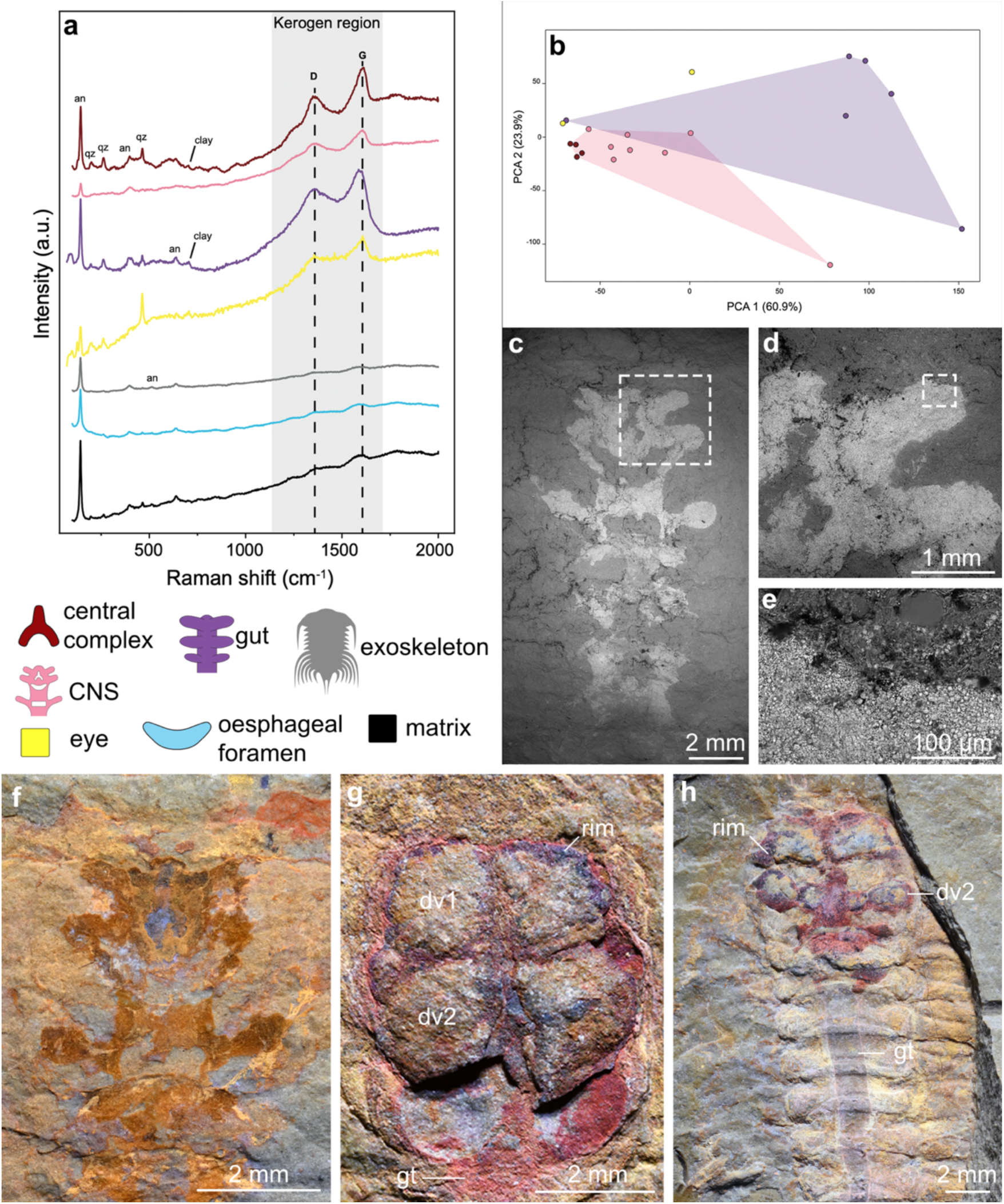
Comparative chemical analyses of nervous and digestive system preservation in *Anacheirurus* from the Fezouata Shale. **a,** Raman spectra from MCZ:IP:202503 and MCZ:IP:202504; see fig. S8 for spot locations. **b,** PCA analysis of Raman spectra from eye and gut of MCZ:IP:202504 compared with the CNS and central complex of MCZ:IP:202503. Panels a and b follow same color scheme shown below panel a. **c – e,** backscattered electron micrographs of nervous system in MCZ:IP:202503 showing framboidal iron deposition. **f,** Nervous system preservation in MCZ:IP:202503. **g,** Cephalic gut diverticulae preservation in *Anacheirurus* sp. (MCZ:IP:202504). **h,** Cephalic gut diverticulae and gut tract preservation in *Anacheirurus adserai* (MCZ:IP:202508). Institutional abbreviations: Museum of Comparative Zoology, Harvard University (MCZ). Abbreviations: an, anatase; dv, diverticula; gt, gut tract; rim, rim of iron; qz, quartz.

MCZ:IP:202503 represents part and counterpart of the anterior half of an adult individual with a length of 17.2 mm and maximum cephalic width of 13.5 mm preserved in dorsoventral view and showing clear evidence of non-biomineralized tissues including the uniramous antennae and internal organs (Fig. 1b, c; fig. S1). The internal anatomy of MCZ:IP:202503 is expressed as a compressed bilaterally symmetrical darkened axial structure with a complex organization that spans almost the entire length of the specimen but does not reach the anterior cephalic edge. The axial structure’s anterior margin has a bulbous median protrusion, followed posteriorly by a metameric series of laterally splayed lobes. In the cephalic region, the axial structure has three lobes that become progressively larger towards the back of the head, followed by a further three narrower and evenly spaced lobes located in the thorax (Fig. 1d-f). The axial structure also shows a series of median features, the first one located anteriorly and having a furcula-like shape (see ‘central complex’ in Fig. 1e) with a blueish grey coloration, immediately followed by a larger roughly V-shaped opening located centrally within the cephalon (see ‘esophageal foramen’ in Fig. 1e), and then a series of rectangular-shaped openings more posteriorly associated with the thorax (Fig. 1b, c). The metameric organization and overall morphology of the symmetrical axial structure in MCZ:IP:202503 is consistent with the euarthropod CNS^33,35,37,38,40^, further strengthened by its chemical composition, preservation, and comparisons with other components of the internal anatomy.

### *Anacheirurus* central nervous system preservation

Backscattered electron microscopy combined with energy dispersive spectroscopy and Raman spectroscopy (see Supplementary methods) demonstrate that the fossilized CNS in MCZ:IP:202503b is replicated by the iron oxide goethite, with several regions enriched in macromolecular carbon (Fig. 2a; figs. S8-10, Supplementary Discussion). The selective enrichment of iron (Fig. 1e) and carbon (Fig. S2) in the CNS contrasts with the clay-rich composition of the exoskeleton and surrounding matrix, which are characterized by higher intensities of aluminum (Fig. 1f), magnesium and potassium (Fig. S2). Backscattered scanning electron microscopy reveals that the iron oxides replicating the CNS are framboidal pyrite pseudomorphs (Fig. 2c-e). The presence of framboids clearly restricted to the CNS support its preservation via the anaerobic decay of organic matter by sulfate-reducing bacteria during early diagenesis. This mode of preservation is comparable to reports of neurological remains of early Cambrian euarthropods such as *Fuxianhuia protensa*^33,34^ and *Alalcomenaeus* sp.^35^, and supports the hypothesis that pyritization of the nervous system was facilitated by their labile nature and originally high concentrations of biogenic iron in the form of ferritin^48,49^. This is also in accordance with the preservation pathways previously proposed for the Fezouata biota, where less recalcitrant tissues were three-dimensionally preserved via pyritization, while decay-resistant tissues were originally preserved as carbonaceous compressions^48,49^. However, the extensive weathering experienced by the Fezouata Shale is largely responsible for leaching most of the carbonaceous matter from the fossils in the site^48^.

Elemental mapping shows that the furcula-shaped feature between the first and second set of paired lobes is depleted in iron relative to the surrounding CNS (Fig. 1e) but still exhibits higher iron intensities than the more posterior V-shaped opening or the exoskeleton (Figs. S4 – 6). Critically, the furcula-shaped feature is visually distinct from the CNS and V-shaped opening under light photography, exhibiting a distinctive dark grey to blueish hue in both part and counterpart (Fig. 1b, c; fig. S1). Raman spectroscopy confirms the presence of organic carbonaceous matter in the furcula-shaped feature as demonstrated by the characteristic D and G bands, whereas it is absent or only detected in very low levels in the V-shaped opening, exoskeleton and rock matrix (Fig. 2a). The presence of carbonaceous remains explains the distinctive coloration of the furcula-shaped feature and support its interpretation as part of the CNS, whereas the V-shaped opening is best explained as an orifice (Fig. 2a). Principal component analysis of several Raman parameters, derived from deconvolution of the D and G bands from MCZ:IP:202503 and MCZ:IP:202507, shows that the furcula-shaped structure groups with the CNS (Fig. 2b). In contrast, spectral features from the digestive system of MCZ:IP:202507 occupy distinct regions in the PCA plot the CNS of MCZ:IP:202503 (Fig. 2b and Fig. S10 and Supplementary Discussion); these discrepancies most likely stem from the differences in the original histological composition between the fossilized internal organs.

### *Anacheirurus* digestive system morphology and preservation

New data on the digestive tract of *Anacheirurus adserai* further strengthens our identification of neurological remains in MCZ:IP:202503 (Fig. 3b). The cephalon of *A. adserai* features three-dimensionally preserved paired gut diverticulae associated with the glabella (Figs. 2g, h, 3f). The cephalic gut diverticulae consist of five simple lobe-like lateral extensions of the digestive tract of almost even size and spacing, and which are accommodated between the glabellar furrows (Fig. 3a, c). The anteriormost pair of diverticulae occupies the frontal lobe, and each succeeding pair aligns with the glabellar lobes (Fig. 3e). Their convex appearance results from early diagenetic mineralization, most likely microbially mediated phosphatization and pyritization^14^, evidenced by pseudoframboids (Fig. 1e; Fig. S3c) and occasional detection of the v1 PO₄³⁻ (964 cm⁻¹) vibrational mode found in apatite (Fig. S7 and Supplementary Discussion), which is completely absent from the CNS. Therefore, the cephalic gut diverticulae in the studied material of *A. adserai* are morphologically, compositionally, and taphonomically distinct from the bilaterally symmetrical axial structure corresponding to the CNS. The cephalic gut diverticulae further differ from the CNS in their size and spacing, lack of medial structures (i.e. furcula- and V-shaped orifice), retain a convex profile (Fig. 1, 3c, d; Fig. S1), and exhibit spectroscopic signatures that likely reflect distinct original compositions and/or degradation pathways (Fig. 2b and Fig. S9 and Supplementary Discussion). These represent multiple independent lines of evidence that the bilaterally symmetrical axial structure preserved in MCZ:IP:202503 corresponds to legitimate remains of the fossilized CNS, rather than preservation artefacts of the digestive tract or other components of the internal anatomy.

**Figure 3.**
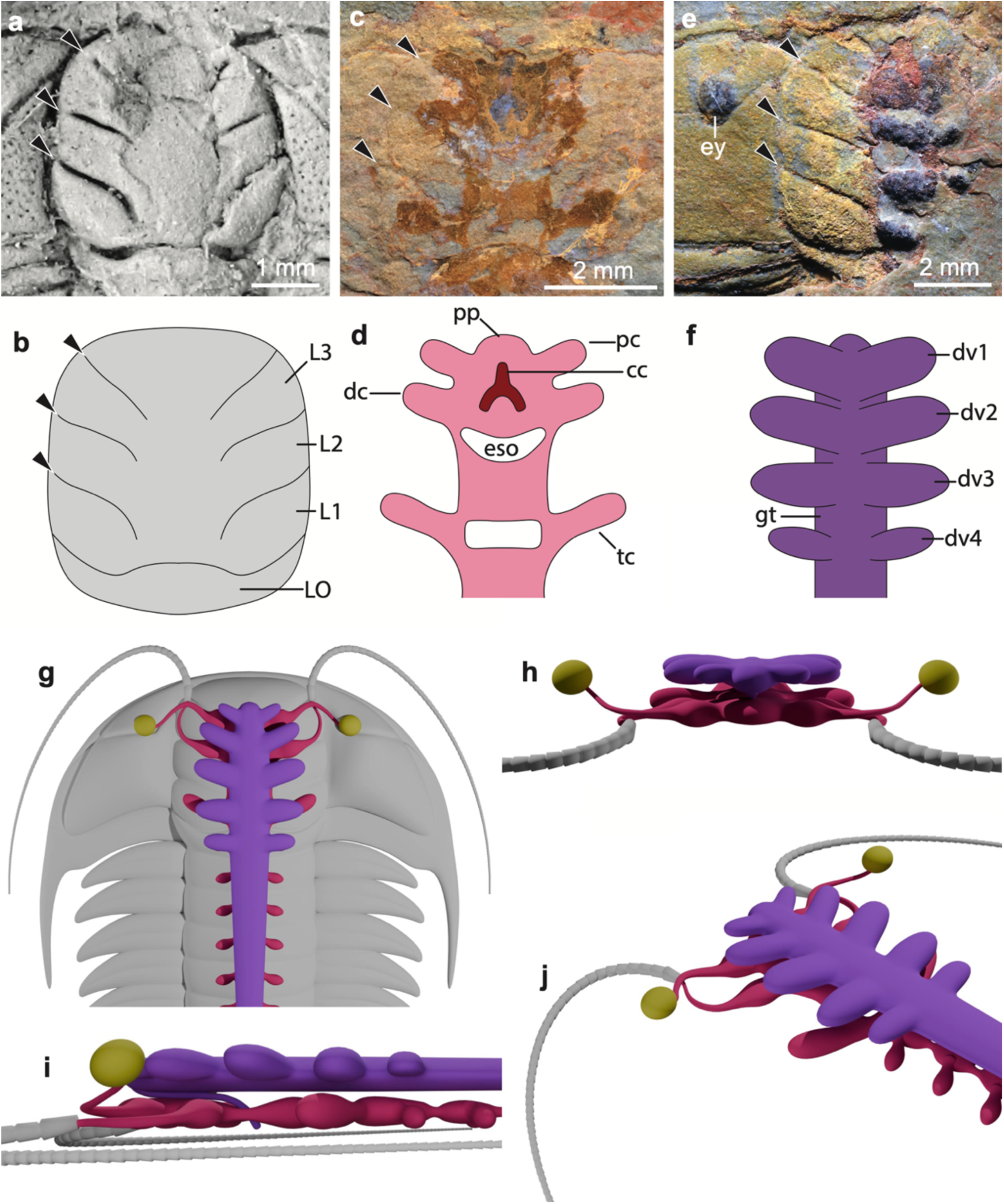
Morphological comparison and reconstructions of cephalic external and internal anatomy in the trilobite *Anacheirurus adserai* from the Fezouata Shale. **a,** ML20-269198, glabella; arrowheads indicate position of glabellar furrows for comparative purposes. **b,** Schematic diagram of glabella. **c,** MCZ:IP:202503, central nervous system. **d,** Schematic diagram of central nervous system. **e,** MCZ:IP:202507, gut diverticulae. **f,** Schematic diagram of gut diverticulae. **g-j,** Three dimensional morphological reconstructions of internal anatomy of *Anacheirurus adserai* by Franz Anthony. **g,** Dorsal view. **h,** Anterior view. **i,** Lateral view. **j,** Oblique view of central nervous system. Institutional abbreviations: Museum of Comparative Zoology, Harvard University (MCZ); Natural History Museum of Lyon, France (ML). Abbreviations: cc, central complex; dc, deutocerebrum; dv*n*, gut diverticula; gt, gut tract; eso, esophageal foramen; L*n*, glabella lobe; LO, lateral occipital lobe; pc, protocerebrum; pp, protocerebral protrusion; tc, tritocerebrum.

## DISCUSSION

### Taphonomy of MCZ:IP:202503

MCZ:IP:202503 does not show a direct connection between the anterior brain neuromeres with the eyes (i.e. optic nerve) and the uniramous antennae, which differs from most instances of cephalic CNS preservation in Cambrian euarthropods^33,37,40^. We posit that this condition can be explained by the fact that *A. adserai* represents the first biomineralizing euarthropod for which a fossil CNS has been reported so far, and the dorsal position of the sessile compound eyes of trilobites. All other instances of euarthropod CNS preservation to date have been reported from non-biomineralized representatives with stalked eyes (i.e. radiodonts^39^, fuxianhuiids^33,34,43^, megacheirans^35,38,42^, hymenocarines^37^, mollisoniids^44^), which facilitates the identification of delicate neurological tracts such as the optic nerve through the cuticle. The presence of a biomineralized dorsal exoskeleton would obscure these fine neuroanatomical details unless the specimen were split in precisely the appropriate plane hosting them. Due to the three-dimensional organization of the cephalic CNS in euarthropods, the fact that the splitting plane of MCZ:IP:202503 shows a frontal section of the specimen makes it impossible to observe neurological structures that would be positioned above (e.g. optic nerve) or below (antennal nerves) relative to the brain. In the context of trilobite cephalic anatomy, it is considered that the optic nerve is accommodated within the eye ridge that connects the sessile dorsal eyes with the glabella^50^, and various studies demonstrate that the uniramous antennae attach to the underside of the cephalon at either side of the ventral hypostome^17,51^. The combination of the biomineralized dorsal exoskeleton and the three-dimensional organization of the trilobite head, including sessile compound eyes and ventral antennae, posit a unique context for the correct interpretation of the fossilized morphology in MCZ:IP:202503. This conclusion is further supported by the CNS preservation in the Carboniferous xiphosuran *Euproops danae*^52^, in which the entire prosomal synganglion is clearly expressed but without traces of the nerve tracts that would connect to the dorsal sessile eyes or the ventral appendages.

### General organization of the trilobite central nervous system

The CNS of *Anacheirurus adserai* MCZ:IP:202503 (Fig. 3g-j) most closely resembles that of Cambrian and extant total-group mandibulates^33,34,36,37,40,45,53–56^ (Fig. 4 and fig. S11), from which we can draw several anatomical correspondences. The overall organization of the cephalic region broadly reflects a tri-partite brain, in which each of the progressively developed paired lobes represent the protocerebral (ocular), deutocerebral (antennal), and tritocerebral (post-antennal) neuromeres respectively (Fig. 1e; 3). The central location and size of the conspicuous V-shaped opening is consistent with the esophageal foramen that accommodates the posteroventral-facing J-shaped foregut in extant euarthropods^45^. Indeed, the location of the esophageal foramen at the level of the second lobe of the glabella in *A. adserai* (Fig 3a-d) matches recent three-dimensional data illustrating the antero-ventral curvature of the trilobite foregut^6,51^. The position of the paired neuromeres relative to the esophageal foramen directly matches their segmental identity, with the first (protocerebral/ocular) in a pre-oral position, the second (deutocerebral/antennal) in a para-oral position, and the third (tritocerebral/post-antennal) in a post-oral position^15–17^. The narrower paired lobes separated by rectangular openings located in the thorax of MCZ:IP:202503 correspond to the ventral nerve cord’s condensed ganglia linked transversely by commissures and longitudinally by connectives (Fig. 1e), as widely expressed in extant and Cambrian euarthropods^43,44,56,57^.

**Figure 4.**
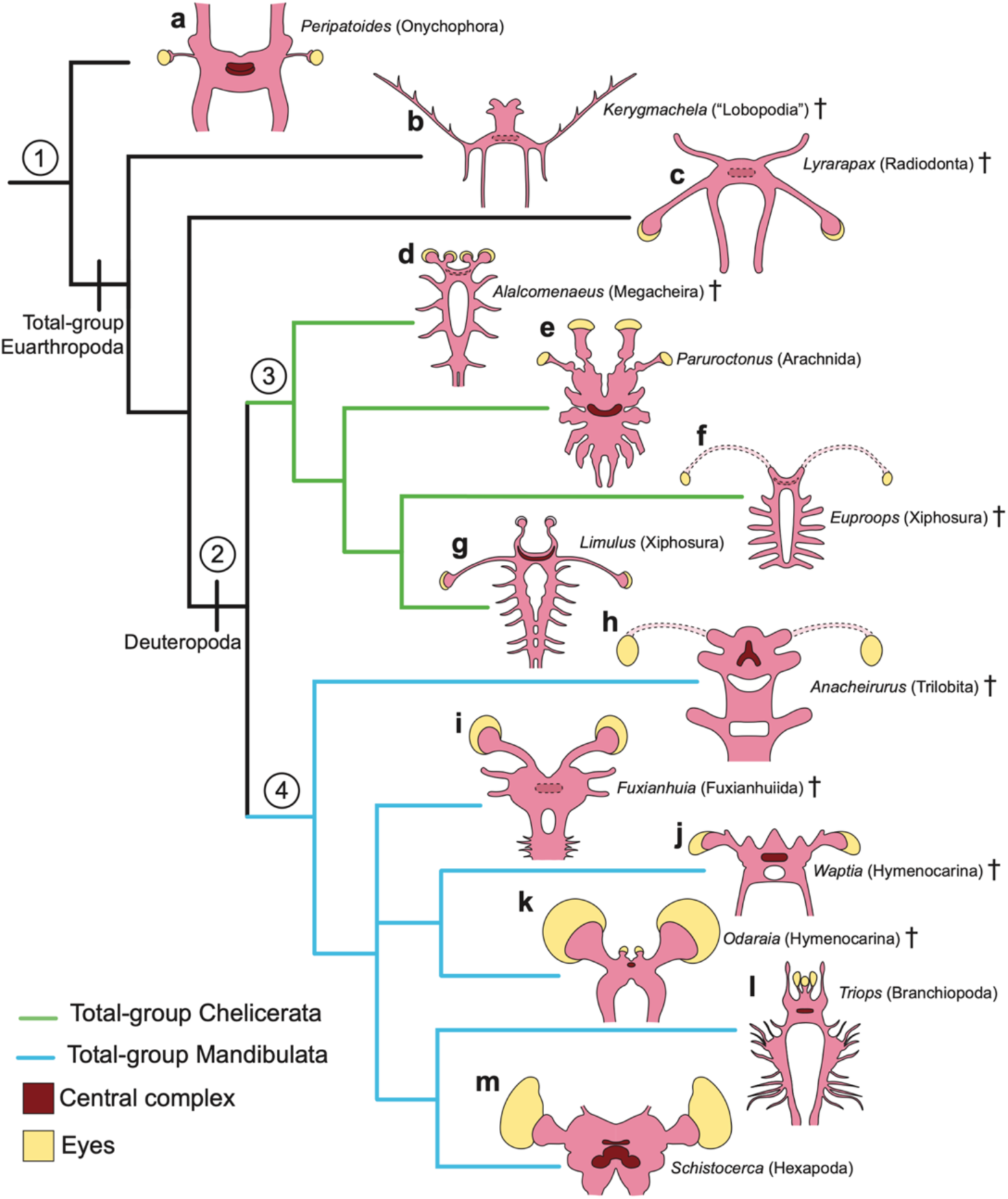
Central nervous system macroevolution in extant and extinct panarthropods. **a,** The extant onychophoran *Peripatoides leuckarti* (redrawn from^54^). **b,** The Cambrian lower stem-group euarthropod *Kerygmachela* (redrawn from^41^). **c,** The Cambrian radiodont *Lyrarapax* (redrawn from^44^ based on^39^). **d,** The Cambrian megacheiran *Alalcomenaeus* (redrawn from^38^). **e,** The extant scorpion *Paruroctonus* (redrawn from^45^). **f,** The Carboniferous xiphosuran *Euproops* (modified from^52^). **g,** The extant xiphosuran *Limulus* (redrawn from^44^, based on^76^). **h,** The Ordovician trilobite *Anacheirurus*. **i,** The Cambrian fuxianhuiid *Fuxianhuia* (redrawn from^44^ based on ^33,43^). **j,** The Cambrian hymenocarine *Waptia* (redrawn from^37^). **k,** The Cambrian hymenocarine *Odaraia* (redrawn from ^40^). **l,** The extant branchiopod *Triops* (redrawn from ^46^ based on ^77^). **m,** The extant orthopteran *Schistocerca* (redrawn from ^54,71^). **1.** Central complex consists of compact laminated midline neuropil (e.g. onychophoran “arcuate body”). **2.** Condensed tri-partite brain with deutocerebral first appendage pair. **3.** Central complex bent anteriorly (i.e. chelicerate “arcuate body”), and deutocerebral raptorial appendage / chelicerae. **4.** Central complex with protocerebral bridge and lateral accessory lobes, and deutocerebral flagelliform first antennae. Central complex diversity in extant mandibulates demonstrates numerous instances of secondary simplification^45,53,55^. Dashed lines denote neurological structures that have not been directly observed in fossil taxa.

The neuromeres in MCZ:IP:202503 would innervate the dorsal compound eyes, the uniramous antennae, and the biramous first appendage pair respectively in relation to *A. adserai*’s external anatomy^46^. Although this general neurological organization is typical for extant euarthropods, MCZ:IP:202503 suggests that the trilobite CNS had some unique attributes (Fig. 3). Most significantly, the fossilized nervous system shows a closer integration between the paired lobes corresponding to the protocerebral/ocular and deutocerebral/antennal neuromeres in the brain, whereas the tritocerebral/post-antennal neuromere occupies a more distant position posteriorly. This precise arrangement deviates from that of the highly condensed brain of extant euarthropods, in which the three brain neuromeres are fused together into a compact syncerebrum. This difference could be regarded as evidence that the trilobite brain consisted of only two fully integrated neuromeres (i.e. bipartite), and thus capture an ancestral organization compared to the tri-partite brain of crown-group euarthropods. However, another possible interpretation is that the brain of *A. adserai* reflects the low degree of functional tagmosis that characterizes the appendages of the trilobite head. Other than the uniramous antennae, the cephalic limbs of trilobites consist of largely homonomous biramous appendages (see ^20^), with only rare known instances of leg reduction for the first pair of post-antennal appendages^51^. Trilobite feeding employed a low degree of limb differentiation when contrasted with the complex feeding capabilities observed in the gnathal appendages of extant euarthropods, which incorporate a combination of food capture, mastication and manipulation. These observations suggest that the conspicuous separation of the tritocerebral/post-antennal neuromere expressed in *A. adserai* could reflect the low degree of functional and morphological integration that typifies the trilobite head. The observed distance between the brain neuromeres in *A. adserai* is likely compounded by the fact that trilobites have proportionally larger cephalic tagma relative to their overall body size when compared with most crown-group euarthropods, resulting in a more outstretched anterior CNS. The close morphological resemblance between the tritocerebral/post-antennal neuromere and the paired lobes in the thorax of MCZ:IP:202503 suggest that the evolution of a fully integrated tri-partite brain involved the co-option of the anterior ventral nerve cord. A comparable example may be found in extant onychophorans (velvet worms), in which the slime-papillae correspond to the third (tritocerebral) head segment in terms of their relative position and function, but their posterior location and innervation is most closely comparable with that of the ventral nerve cord (e.g. ^58^).

### Significance of the anteromedian protocerebral protrusion

The anteromedian protocerebral protrusion in MCZ:IP:202503 resembles the protocerebral innervation of the median eyes observed in branchiopod crustaceans in terms of their morphology and position^56^. A similar anteromedian protrusion has also been reported in the Burgess Shale hymenocarines *Waptia fieldensis*^37^ and *Odaraia alata*^40^ associated with the so-called anterior sclerite that has a protocerebral segmental origin (Fig. S11 and Supplementary Discussion). Trilobites lack a discrete anterior sclerite, but this structure has a deep evolutionary origin and is widespread among non-biomineralized trilobitomorphs, fuxianhuiids, bivalved forms, and other Cambrian euarthropods^40^, but have median eyes located on the glabella ^59^. We propose that the anteromedian protocerebral protrusion in MCZ:IP:202503 might reflect the ancestral innervation between the CNS and the anterior sclerite following the recent hypothesis that this exoskeletal plate became incorporated into the trilobite hypostome^60^.

### The central complex of *Anacheirurus adserai*

The furcula-shaped feature observed in MCZ:IP:202503 is notable in terms of its intricate morphology and position on the anterior CNS of *Anacheirurus adserai* (Fig. 1e, h). Comparisons with the brain of extant euarthropods show that the position and shape of the furcula-like feature closely match the outline of the central complex, a highly conserved region of the protocerebrum that consists of a dense layered cluster of midline neuropils that plays a key role in vision, navigation and motor control^45,53–55,57^. Transverse sections of the mandibulate central complex in dorsal view illustrate that the morphological continuity between the protocerebral bridge, the central body, and the paired lateral accessory lobes (Fig. 1g) convey the distinctive furcula-like outline observed in MCZ:IP:202503 (Fig. 1e, h), although the precise shape and components of the central complex can differ substantially across extant mandibulates^54,55^. The CNS remains preserved in MCZ:IP:202503 are evenly distributed between both part and counterpart as evidenced by both light photography (Fig. 1b, c; fig. S1) and elemental mapping (fig. S4-S7), which indicates that this specimen effectively captures a frontal section of the fossilized internal anatomy (i.e. dividing the body into an upper and lower parts). This is significant because the central complex of euarthropods is embedded within the condensed dorsal brain^53–55^, and thus would only be observable through a cross-sectional view of the CNS. Furthermore, the differences observed between the furcula-shaped feature (i.e. dark grey to bluish coloration, stronger carbonaceous signal, lower iron intensity) and the rest of the surrounding CNS (i.e. orange coloration, comparatively weaker macromolecular carbon signal, high iron intensities) could be explained by discrepancies in the original histology of these structures. Given that the central complex is a midline neuropil, it consists of a dense array of axons, dendrites and glial cells embedded within the brain, whereas the cell bodies are located on the periphery of the brain, creating compositional heterogeneity with a higher concentration of organelles and other intracellular elements on the outer layers of the CNS. We hypothesize that the more pristine carbonaceous preservation of the central complex in MCZ:IP:202503, as indicated by a stronger carbonaceous signal and less pyritization, could have resulted from its deeper position within the brain and fortuitous split plane, whereas the replication of the majority of the CNS through pyritization would have been driven by the consumption of the organelle-rich peripheral cell bodies by sulphate reducing bacteria.

### Significance of the trilobite central complex

The general shape of the central complex is of critical phylogenetic significance because the furcula-like outline suggests the presence of a protocerebral bridge and lateral accessory lobes, which are exclusive to mandibulates, and specifically some pancrustaceans including hexapods, in the context of panarthropod CNS diversity^45,53–55,57^. By contrast, the corresponding midline neuropil found in the brain of onychophorans and chelicerates, also known as the arcuate body, has instead a distinctly flattened or anteriorly bent crescent shape without lateral accessory lobes or a distinct protocerbral bridge^54^ (Fig. 4a, g). Albeit simple, the overall outline morphology of the central complex and its constituent structures are characters that fundamentally discriminate the neuroanatomy of the two major groups of living euarthropods^57,61,62^. It should also be noted that the precise organization of the central complex in Mandibulata is highly variable, and most likely involves several instances of convergent secondary loss or simplification of some of its components. Within Pancrustacea there is patchy distribution of midline neuropil characters in different groups. Whereas the central complex of eumalacostracans and hexapods typically feature a well-developed layered central body (composed of the ellipsoid body and fan-shaped body) and a protocerebral bridge, some of these features are comparatively reduced in other clades such as Ostracoda, Copepoda, Branchiopoda and Remipedia^55,57,63^ (Fig. 5). More broadly, the central body of Chilopoda resembles the arcuate body of Chelicerata in the presence of a simple ellipsoid body, whereas a central complex appears to be absent in Diplopoda^57^ or extremely simple^63^. In this context, the central complex of *Anacheirurus adserai* is incompatible with the arcuate body of onychophorans and chelicerates, and instead more readily comparable with the central complex of eumalacostracans and hexapods. The paucity of neurological information for Myriapoda, coupled with their seemingly simplified organization, create some uncertainty when attempting to polarize the evolution of the euarthropod central complex (see discussion below). A central complex has not been formally recognized in the fossilized CNS of extinct euarthropods so far. However, topological comparisons with the fossilized CNS of the Burgess Shale *Waptia fieldensis*^37^ and *Odaraia alata*^40^ suggest that enigmatic anteromedian features in the brain of these taxa might also correspond to potential remains of the central complex (Fig. S11 and Supplementary Discussion).

**Figure 5.**
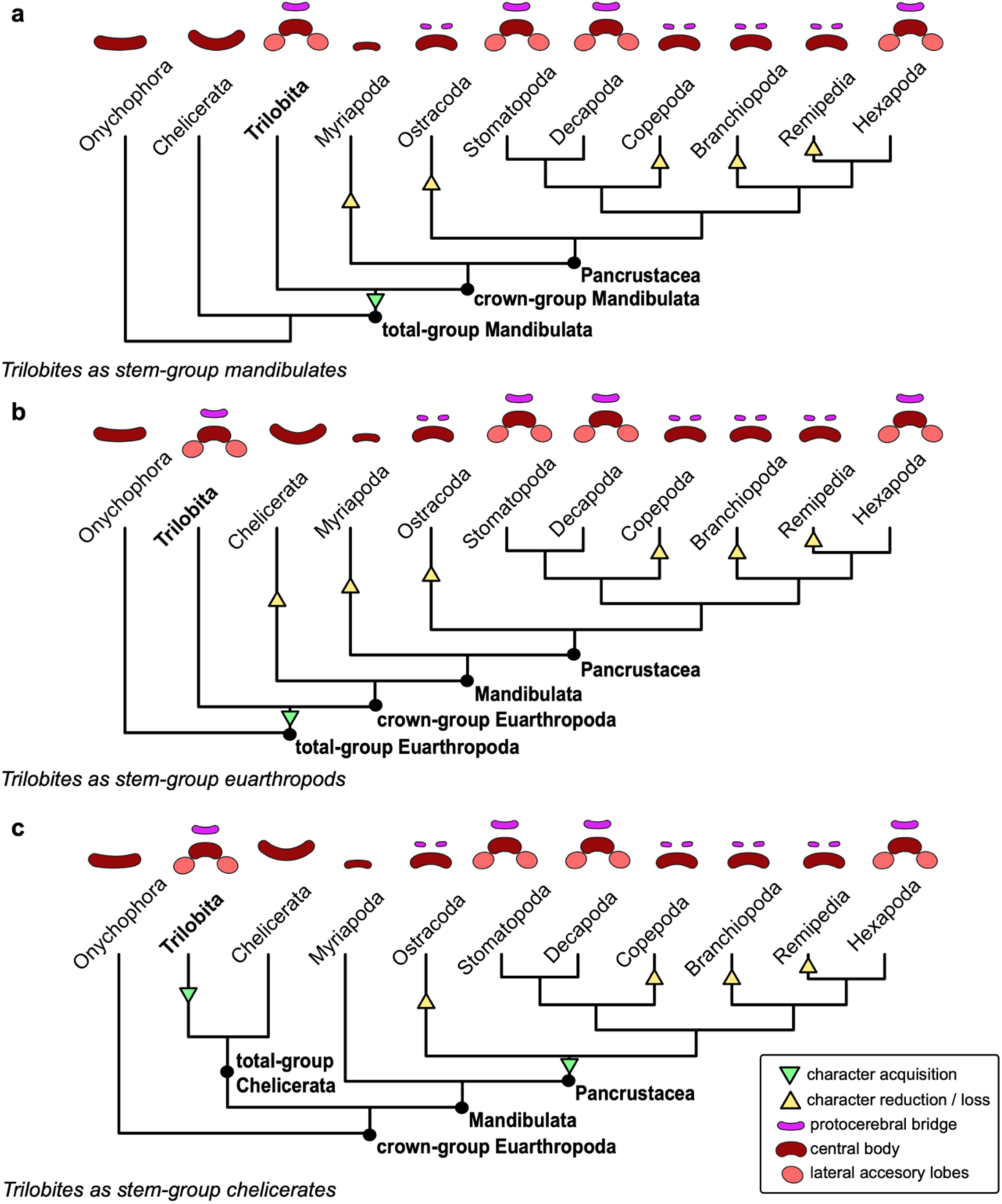
Competing implications of central complex morphology in trilobite *Anacheirurus adserai* for euarthropod evolution. **a.** Trilobites as stem-group mandibulates; the scenario suggests repeated secondary reduction or loss of central complex characters within Mandibulata, including myriapods, ostracods, copepods, branchiopods and remipedes. **b.** Trilobites as stem-group euarthropods; the scenario suggest a deeper origin for the central complex within the euarthropod stem-lineage, with the presence of an arcuate body in onychophorans and chelicerates resulting from the loss of characters in the latter lineage. **c.** Trilobites as stem-group chelicerates; the scenario suggests the convergent evolution between the central complex of trilobites and pancrustaceans (depicted), or alternatively the origin of the central complex in the chelicerate stem-lineage and secondary character loss in crown-group Chelicerata (not shown in diagram). Overall topology and schematic diagrams of central complex diversity modified from refs ^55,63^.

### Trilobites as early mandibulates

The CNS of *Anacheirurus adserai* shares several derived characters with Cambrian^33,34,36,37,40,45^ and extant mandibulates^53–57^ (Fig. 4). Our paleoneuroanatomical data suggests that trilobites belong to stem-group Mandibulata (Fig. 4h), whereas the paucity of synapomorphic characters in terms of their exoskeletal and appendicular morphology, such as the absence of mandibles or other specialized gnathal appendages, argues against a position within the crown-group. Our findings also allow us to better contextualize prior evidence for mandibulate affinities. The discovery of crystalline cones and rhabdoms in the compound eyes of Cambrian, Ordovician and Devonian trilobites led to the interpretation that they reflect the ancestral organization of the mandibulate compound eye^21–23^, here further supported by the CNS organization. Our results also strengthen the hypothesis that the trilobite uniramous antennae are homologous to the deutocerebral first antennae (or antennule) of mandibulates^15–17^, and suggest that the first biramous appendage pair of trilobites is serially homologous with the tritocerebral second antennae of non-hexapod pancrustaceans. This segmental alignment carries the implication that the ancestral mandibulate tritocerebral appendage pair had a biramous organization and was used primarily for locomotion and feeding, as observed in most trilobites, and that the sensorial function observed in pancrustaceans represents a derived condition. Notably, exceptional three-dimensional data from early Cambrian trilobites^51^ and non-biomineralized trilobitomorphs^64^ show evidence for the secondary modification of the tritocerebral appendage for a sensorial function, including a reduction of the endopodite and filiform morphology of the exopodite; these instances may capture early stages of the evolution of sensorial tritocerebral appendages in the mandibulate lineage. Given that the main support for chelicerate affinities for trilobites comes from the overall similarity of the dorsal exoskeletons and comparisons with the ventral ancestrally biramous appendages relative to xiphosurans^5,7,13^, we conclude that these represent analogous functional and ecological adaptations to a similar mode of life in the benthic marine environment^65^. The position of trilobites also impacts the overall species composition of the mandibulate stem lineage. Trilobites are part of the morphologically disparate clade Trilobitomorpha that includes numerous non-biomineralized taxa united by the presence of a broad dorsal exoskeleton, uniramous antennae, biramous appendages with a stenopodous endopodite and lamellae-bearing exopodites^15,16,18^. The affinity of trilobites, and by extension antennae-bearing trilobitomorphs, as stem-group mandibulates weakens the longstanding Arachnomorpha hypothesis that these taxa represent chelicerate ancestors (*contra* ^24,26,66^).

### Early evolution of the euarthropod central complex

The central complex represents a fundamental component of the brain of extant Euarthropoda and Onychophora^45,53,55,57,67^, and thus *Anacheirurus adserai* directly informs the macroevolution of this structure in deep time. The similarities of the central complex shared between onychophorans and chelicerates most likely reflect the ancestral organization for Panarthropoda given their phylogenetic position (see also^45,63^), namely a compact midline layered neuropil with a laterally elongate outline (Fig. 4a), particularly considering that the anteriorly-bent crescent shape is exclusive to chelicerates (Fig. 4e, g). These observations imply that Cambrian lower stem-group euarthropods for which CNS preservation is known, such as the flap-bearing lobopodian *Kerygmachela*^41^ and the radiodonts *Lyrarapax*^39^ and *Stanleycaris*^68^, most likely featured an onychophoran-like layered and flat central complex overall morphology (Fig. 4b, c; Fig. 5). Consequently, it appears most likely that the known CNS of Cambrian euarthropods regarded as stem-group chelicerates, namely megacheirans^35,38^ and mollisoniids^44^, would have also featured an arcuate body comparable with that of extant chelicerates^67^. The new information on the neuroanatomical organization of *A. adserai*, coupled with the controversial phylogenetic position of trilobites, results in three major scenarios for the evolution of the euarthropod central complex. We discuss these in turn, from most to least likely in our view given the available data.

### Trilobites as early mandibulates

The CNS of *A. adserai* suggest that trilobites had a central complex featuring a protocerebral bridge and lateral accessory lobes, making it most comparable to that of some extant pancrustaceans including hexapods. The position of trilobites in stem-group Mandibulata, that is branching off before the divergence between Myriapoda and Pancrustacea, would imply that the modification of these neuroanatomical features in various groups such as myriapods, ostracods, branchiopods and remipedes^45,57^, have likely experienced various degrees of secondary simplification and/or loss of these characters^45,53,55^ (Fig. 5). This pattern would be particularly significant for myriapods, as the simple neurological organization of the central complex observed in chilopods and diplopods^45,57^ would be best regarded as highly derived for this lineage in particular, rather than ancestral for Mandibulata as a whole. This scenario would in turn indicate that an intricate central complex appeared early during mandibulate history and has subsequently experienced extensive mosaic evolution including instances of dramatic reduction or loss of characters (Fig. 5). Critically, secondary simplification or loss of neurological structures relative to close relatives is commonplace in euarthropod evolutionary history, as inferred from extant euarthropod phylogenies informed by neuroanatomical characters^45,62^. More broadly, the position of trilobites as stem-group mandibulates would also imply that the overall organization of the arcuate body as expressed in onychophorans and chelicerates is closer to the ancestral condition for Panarthropoda, whereas the protocerebral bridge and lateral accessory lobes are innovations of total-group Mandibulata.

### Trilobites as stem-group euarthropods

The alternative position of trilobites as upper stem-group euarthropods would indicate that a pancrustacean-like central complex with a protocerebral bridge and lateral accessory lobes is ancestral for at least part of Deuteropoda^19,69^. In turn, this scenario would imply that the similarities between the arcuate body of onychophorans and chelicerates are convergent as a result of character loss, rather than ancestral as hypothesized based on neontological neuroanatomical information^54,55,63^. In a broader context, this model would carry similar implications for mandibulate evolution to the scenario of trilobites as stem-group mandibulates (above), with the only difference that a pancrustacean-like central complex would have a deeper origin during euarthropod evolution.

### Trilobites as early chelicerates

The third possibility is that trilobites are stem-group chelicerates, as widely considered historically due to their superficial resemblance with horseshoe crabs^24,26,70^. As established before, the similarities between the arcuate body of extant onychophorans and chelicerates suggest that they capture the ancestral organization for the panarthropod central complex. Thus, the CNS organization of *Anacheirurus adserai* would require either a striking case of convergent evolution between trilobites and pancrustaceans, or alternatively the gain and subsequent loss of neurological character within the chelicerate stem-lineage (Fig. 5). Both options are weakly supported in our view and would require a considerable series of unparsimonious evolutionary changes in the CNS for which there is no additional supporting evidence available.

### Trilobite brains and the Cambrian Information Revolution

Trilobites are the stratigraphically oldest euarthropods in the fossil record^1,10,11^, so our findings imply that the ancestral mandibulate CNS and central complex can be traced to the base of the Cambrian Stage 3 (ca. 521 million years ago), assuming that the earliest representatives of this clade had a broadly comparable neuroanatomy. In modern insects, the central complex plays crucial roles in locomotor control, polarized light vision, orientation and navigation^71,72^. As such, the trilobite CNS illuminates the early evolution of complex behaviors during the Cambrian Information Revolution defined by the proliferation of locomotory and feeding traces observed in the Fortunian^30,73,74^. Given the paucity of Terreneuvian bilaterian body fossils or internal anatomical data, the primary evolutionary innovations associated with the Cambrian Information Revolution have been restricted to external features with a sensorial function such as eyes and antennae^15,21–23^. The central complex of *A. adserai* offers the first direct evidence of the neurological adaptations for sophisticated visual prowess, sensorimotor processing and navigation in trilobites responsible for the production of intricate trace fossils as well as spectacular instances of collective and coordinated behavior throughout the Paleozoic Era^75^. Ultimately, trilobite paleoneuroanatomy supports the view that the euarthropod CNS has retained extraordinarily conservative elements for the entirety of the Phanerozoic Era, and suggests that the fundamental neurological diversification and innovation observed among extant Panarthropoda most likely took place within the interval encompassing the latest Ediacaran and the earliest Cambrian ^45^.

## Acknowledgements

The authors thank Jessica Cundiff, Karma Nanglu, and Jared C. Richards (Museum of Comparative Zoology, Cambridge, USA) and Jessica Utrup (Yale Peabody Museum, USA) for processing and facilitating access to specimens, Franz Anthony (https://franzanth.com/) for the morphological reconstruction, Lukáš Laibl (Czech Academy of Sciences, Institute of Geology, Prague, Czech Republic) for discussion of photography, Uwe Homberg (Philipps-Universität, Marburg, Germany) for sharing histological photographs of the locust central complex, and Jean-Bernard Caron (Royal Ontario Museum, Toronto, Canada) for sharing digital images of *Waptia*. We acknowledge the key role of Mohamed ‘Ou Said’ Ben Moula and the Ben Moula Family in the discovery, collection and characterization of the studied fossil material and their contributions towards facilitating access the scientific study of the Fezouata Shale biota of Morocco. Thanks to Allison Daley (University of Lausanne, Switzerland), Peter Van Roy (Ghent University, Belgium) and Stephen Pates (University of Exeter and University College London, UK) for their assistance recording field data. This work was supported by a Research Grant from HFSP (Ref.-No: RGY0056/2022, the NSF CAREER award No. 2047192 “*Ecological turnover at the dawn of the Great Ordovician Biodiversification Event -quantifying the Cambro-Ordovician transition through the lens of exceptional preservation*”, Lemann Brazil Research Fund, and the Museum of Comparative Zoology Biodiversity Postdoctoral Fellowship. We thank Brigitte Schoenemann, Graham Budd, and Pierre Gueriau for their helpful comments to improve this work.

## Author contributions

S.R.L. photographed specimens, performed elemental analyses/mapping and drafted the first version of the figures and manuscript; B. B. K. collected, analyzed and visualized Raman spectra; J. W. collected SEM backscattered images and EDS spectral and mapping data; F.P.-P. performed photography; J.O.-H. designed the research, led neuroanatomical interpretation and performed final manuscript writing/editing. All authors contributed towards research, discussion and approved the final version of the manuscript.

## Supplemental Material

### Materials and Methods

#### Supplemental Text

Figs. S1 to S11

Table S1

Table S2

Table S3

