## Supplemental Information for "The trilobite nervous system and the early evolution of the mandibulate central complex"

### **The PDF file includes:**

Materials and Methods
Supplementary Text
Figs. S1 to S11
References

### **Materials and Methods**

#### Specimen Details

The studied specimens were collected by Mohammed ‘Ou Said’ Ben Moula from the Draa Valley in the Zagora region of Morocco as part of a collection purchased in 2019 from Lahcen Ben Moula and exported by Brahim Tahiri with the approval of the Ministry of Mines in Rabat (Invoice N. 92/E/21). Geolocation of the sub-localities that produced the studied specimens are curated with their respective collections at either the Museum of Comparative Zoology (MCZ) or the Yale Peabody Museum (YPM).

#### Specimen photography:

All specimens were photographed under cross-polarized light using a Nikon D850 DSLR camera with a Macro Nikkor 60 mm lens. MCZ:IP:202503 was photographed submerged in 70% ethanol (diluted with distilled water) to enhance the contrast of the internal anatomy (fig. S1).

#### Scanning Electron Microscopy (SEM) and Energy Dispersive Spectroscopy (EDS)

Three specimens were selected for elemental analyses based on the preservation of the central nervous system, digestive system, and quality of the exoskeleton (fig. S2). Specimens were analyzed at Williams College using a Thermo Scientific Quattro S scanning electron microscope (SEM) fitted with an Everhart Thornley, concentric backscatter, and EDAX Octane Elect detectors. Energy dispersive spectroscopy (EDS) spot analyses (30 second dwell) and mapping, and backscattered electron (BSE) conducted in a high vacuum environment under 20kv operating conditions (see fig. S3 for spot locations). Additional backscattered SEM imaging and EDS elemental mapping were performed on MCZ:IP:202503 (in its native state) using a custom Tescan

Vega GMU (20 kV, WD: 15 mm) variable pressure scanning electron microscope equipped with two Bruker XFlash 5030 x-ray detectors at Harvard University (fig. S4). Since large-scale analysis of the fossil specimen exhibited distinct regio-specific elemental distributions (fig. S5), we subsequently performed a more detailed elemental analysis on the different regions of interest (nervous system, central complex, esophageal foramen, and the exoskeleton) with a focus on the Fe EDS peak. To test for more localized feature-specific Fe concentrations, we collected three EDS point spectra from each of the above regions (fig. S6a), the results of which are shown in fig. S6b. While these Fe concentration differences were subtle, they exhibited clear trends, which were supported by subsequently extracted EDS spectra collected from each mapped region of interest (figs. S6c, d, and S7). For these figures, specific regions of interest from each map were selected (figs. S6c, and S7a, c), their constituent EDS spectra were collected and averaged, and subsequently plotted to demonstrate regional trends in Fe concentration (fig. S6d and S7b, d).

#### Raman Spectroscopy

Raman Spectroscopy (RS) was performed using a Horiba LabRam Evolution available at the Center for Nanoscale Systems, Harvard University (see fig. S8 for spot locations). We used a green laser (532nm), a Olympus LMPlanFL N 50x/0.50 objective, 5-10s of acquisition time, 1-3 accumulations, a grating of 1800 gr/mm, a CCD detector, and a hole size of 200-300 $\mu\text{m}$. The spectra were acquired in the 100-2000 $\text{cm}^{-1}$ range. The final laser power at the sample was 1.6-2 mW as measured by a Thorlabs S170C sensor coupled to a PM100D energy meter console.

The spectra were processed in fityk 1.3.1 software. Our pipeline for the analysis of the macromolecular carbon (so-called D and G) bands consisted of: [i] identifying the peaks present in the spectra to make sure that hematite was not present in the analyzed spot; [ii] cutting the spectra in the 900-2000 $\text{cm}^{-1}$ region; [iii] performing baseline correction (cubic spline) by setting fixed points in the 900-1100 and 1700-2000 $\text{cm}^{-1}$ ranges; and [iv] fitting the spectra with pseudo-Voigt line functions with three different procedures (see below). These procedures resulted in several Raman **band** parameters, including band center, height, full width at half maximum (FWHM), area, ratio of band heights and band area, and Raman Band Separation (RBS). The Principal Component Analysis (PCA) was performed in the Past 4.17c software (Hammer et al. 2001).

#### Raman Data Fitting

Fitting approaches can vary substantially across studies, so we employed three distinct fitting methods to avoid or minimize potential biases (see below and fig. S10). The automated pipeline from Henry et al. (2018) could not be used, as the ‘saddle’ was higher than the D-band FWHM in several gut-derived spectra.

#### Unconstrained fitting

This fitting procedure is the least restrictive, and included 4 to 6 bands, selected based on the shape of the D and G regions. Particularly, the presence of shoulders and elevated residuals were used to infer the presence of additional bands. Depending on the spectrum, the fitted bands included the D4, D1, D6, D3, G, and D2 (See Henry et al. 2019 for an explanation of band nomenclature). Fitting is then performed with the minimum number of bands necessary for a good fit. The presence of the D5 band was not evident by visual inspection or from residuals. While it may be argued that D5 could be concealed due to overlap with D4 and D1, we find no compelling evidence to support its inclusion (fig. S10a-c).

#### Fitting G' of Kouketsu et al. (2014)

In this case, the kerogen bands are fitted with four pseudo-Voigt line functions, with the D4 and D3 fixed at $1245\text{ cm}^{-1}$ and $1510\text{ cm}^{-1}$, respectively. Among all the fitting procedures from Kouketsu et al. (2014), this was the one that better suited the spectra across all the datasets (fig. S10d-f).

#### Minimal-component fitting

The third fitting procedure comprises a more conservative approach where the D and G regions are fitted with only two pseudo-Voigt line functions (fig. S10g-i).

### **Supplementary Text**

#### Preservation

##### *Elemental composition*

The exoskeleton elemental composition of the three analyzed specimens of *Anacheirurus adserai* (MCZ:IP:202503, MCZ:IP:202504, and MCZ:IP:202505) is similar, being composed primarily of aluminum, silicon, magnesium and calcium (figs. S2; Table S2). The matrix of MCZ:IP:202503 has an overlapping range of silicon with the exoskeleton, but with a broader range of carbon wt% values for some locations (Table S2). The spots within the matrix of MCZ:IP:202503 have the highest carbon wt% values (Table S2). In MCZ:IP:202505, the surface of glabellae shows the preservation of gut diverticulae composed of $35.56 - 84.02\text{ wt\%}$ iron (Table S2). Iron concentrations within the CNS of MCZ:IP:202503 vary from $25.28 - 51.59\text{ wt\%}$, while carbon has a range of $2.37 - 19.33\text{ wt\%}$ (Table S2). Aluminum ($5.87 - 9.23\text{ wt\%}$), silicon ($6.29 - 9.07\text{ wt\%}$), magnesium ($0.41 - 2.06\text{ wt\%}$) and calcium ($1.08 - 1.59\text{ wt\%}$) are also present in the CNS, but at low intensities (Table S2). Elemental mapping shows the nervous system is enriched in iron and carbon (fig. S2). The furcula-shaped central complex of MCZ:IP:202503 is compositionally similar to the exoskeleton, being primarily composed of silicon and aluminum (Table S2), but is more enriched in iron than either the esophageal foramen or exoskeleton (figs. S6 and S7). The diverticulae and the gut tract of MCZ:IP:202504 have overlapping iron and carbon wt% (Table S2).

Iron is present in the matrix, exoskeleton, digestive system, and nervous system (Table S2), but with different crystal textures indicating different conditions in which it formed (fig. S2c – j). The CNS of MCZ:IP:202503 is preserved as a dark layer composed of iron framboids (fig. S2c, d). Cephalic gut diverticulae are preserved as convex dome or lobe-like structures with iron framboids concentrated around the margins (fig. S2e, f). The center of the diverticulae is similar in composition to the matrix and exoskeleton. Although framboids occur in both the CNS and digestive system, their distribution varies which allows distinguishing between the structures. The CNS is composed entirely of a uniform patina of framboidal iron, whereas the diverticulae are primary Al and Si, with only rims of framboids, seen in light photographs as an orange rim around the outline (fig. S2f). Unlike the CNS and diverticulae, the lobe like structures in the glabella of MCZ:IP:202505 is composed of iron minerals with euhedral habits (fig. S2g), indicating a different formation pathway from either of the other structures.

Given the analytical limitations of elemental mapping alone, particularly in regard to light elements such as carbon, we further investigated the chemical composition of the internal organs using Raman spectroscopy. The furcula-shaped structure is dark in color, unlike the more posterior features (Fig. 1 and fig. S1), but both appear similar in elemental maps (fig. S4). The CNS is enriched in iron and carbon (figs. S2a and S4). The furcula-shaped structure has higher iron intensity than the V-shaped opening (figs. S6 and S7), as well as macromolecular carbon indicated by the characteristic D and G bands (Fig. 2a). Kerogen is an operational definition for macromolecular organic matter that is insoluble in organic solvents. Kerogen is by far the main component of organic matter in fossils, with bitumen (the soluble fraction) constituting a minor component. Although disordered carbonaceous matter can be produced via abiotic processes, such as hydrothermalism, serpentinization and Fischer-Tropsch-type reactions, there is no evidence for these alterations in the Fezouata samples analyzed here (see below). Organic matter is absent or only shows very low intensities in the spectra from the V-shaped opening, exoskeleton and matrix (Fig. 2a). These data supports the interpretation that the furcula-shaped structure was a feature within the brain, united by higher iron intensities and organic macromolecular carbon, whereas the V-shape opening was a whole within the CNS, likely the esophageal foramen (Fig. 2a,b).

##### *Mineral and molecular composition (Raman Spectroscopy, RS)*

RS confirms that the iron oxide preserving the CNS is goethite (α-FeOOH) (fig. 9), as indicated by diagnostic bands at approximately 251, 307, 403, and 688 $\text{cm}^{-1}$ (see De Faria et al., 1997 and De Faria & Lopes, 2007), observed in spectra from the yellowish spots of specimen MCZ:IP:202503 (fig. S9a). Iron (oxyhydr)oxides, specifically goethite (Gth) and hematite (Hm) have been previously detected in soft-bodied euarthropods from the Fezouata (Potin et al., 2023), but for recalcitrant cuticular tissues that likely escaped early pyritization (Saleh et al., 2020). Although hematite is absent from CNS preservation, we interpret the variable Gth/Hm distribution in Fezouata fossils as reflecting telodiagenetic conditions following uplift and weathering, regardless of whether the precursor phases originated from pyrite oxidation or direct authigenic precipitation associated with kerogen oxidation. Similar variations have also been described for

Devonian shelly fossils (Becker-Kerber et al., 2017), and these late-stage transformations are controlled by water availability, pH, and temperature (Cornell and Schwertmann, 2003).

Anatase is consistently detected across all analyzed samples and regions, including the CNS, gut, exoskeleton, and surrounding matrix (e.g., Fig. 2a and fig. S9). This mineral shows characteristic Raman bands at $144\text{ cm}^{-1}$, $\sim 395\text{--}400\text{ cm}^{-1}$ and $636\text{ cm}^{-1}$. The widespread presence of anatase suggests a late diagenetic origin and the low intensity of titanium (0.12 – 1.84 wt%; Table S3) indicate a minor contribution to the mineralogical content of the samples, despite its well-defined and intense Raman peaks (Fig. 2a). This apparent contradiction is explained by the fact that anatase is well known to be a good Raman scatterer (Murad, 1997; Edwards et al. 2006). Apatite was occasionally identified in the gut region of MCZ:IP:202507, as supported by the presence of its main band at $964\text{ cm}^{-1}$.

One of the most significant findings from RS is the presence of carbonaceous signals in the CNS, the furcula-shaped feature, and the gut, confirming the preservation of organic matter in these tissues. This is especially apparent when compared to other tissues and the surrounding matrix (Fig. 2a), where the peaks are weak or absent. The strongest D and G Raman signals were obtained from the darker regions, as observed under reflected light of the microscope (fig. S7). The location of strong carbonaceous matter signals in tissue-specific regions and its virtual absence in nearby regions including host rock argue against abiotic processes that generate disordered carbonaceous material. These would arguably affect the immediate vicinity of the fossils. It is important to note that no veins or fractures are associated with the fossils or their preserved tissues as well. Finally, Saleh *et al.* (2020) have found sedimentary organic matter from fresh core samples of the Fezouata, and maturity parameters based on their Raman spectral features suggest that these rocks were not substantially altered during burial history. These lines of evidence suggest that the organic matter is original from the organism itself and can be classified as organic carbonaceous matter, likely composed mostly of kerogen (i.e., insoluble sedimentary organic matter).

Preliminary visual inspection of the spectra already showed a distinction in the kerogen signals from the CNS and gut tissues (Fig. 2a; fig. S9). This observation is supported by all three spectral fitting approaches. The PCA loadings show that most spectral variance is driven by parameters within the D region (Table S3), particularly the D FWHM in the minimal-component fitting (only D- and G-functions), and the D4 FWHM in the other two fitting approaches (fig. S9; Table S3). Importantly, the consistent tissue separation between CNS and gut regions in the PCA plots across all fitting approaches rules out possible bias in spectra processing. Interestingly, the carbon signals from the eyes of specimen MCZ:IP:202507 do not consistently cluster with those from its gut (Fig. 2b; fig. S9), suggesting that the observed spectral differences are not due to inter-individual or inter-sample variation. Furthermore, the analyzed specimens were collected from spatially close outcrops (pit A16 N 30°30.751', W 5°50.301' and A8 N 30°29.609', W 5°50.887') without significant stratigraphic differences, ruling out thermal maturity as a driver of kerogen variation.

To our knowledge, this is the first study to document compositional differences in kerogen across distinct tissues within a single taxon using Raman spectroscopy. Although we do not attempt to assign specific functional groups to these differences, our findings suggest they result from tissue-specific diagenesis, possibly with mineral influences. Recent studies showed the potential of kerogen parameters for the discrimination of matured and untreated samples from tissues of different biological groups, as well as fossil taxa (Rossi et al. 2024). However, in the case of those fossils, it remains to be investigated whether the differences reflect original tissue composition, inter-sample variability, different diagenetic pathways or variation in thermal alteration. More importantly, both the presence of kerogen in the CNS and furcula-shaped feature, and their distinctive carbon signatures, argues strongly against interpretations that these structures represent preservational artifacts or voids.

##### Central complex preservation in Cambrian hymenocarines

The discovery of a central complex in *Anacheirurus adserai* (MCZ:IP:202503; Fig. 1; fig. S1) prompts a reevaluation of problematic features on the fossilized CNS of Cambrian euarthropods that share a similar topology and size, namely in the Burgess Shale hymenocarines *Waptia fieldensis* (Vannier et al. 2018) and *Odaraia alata* (Ortega-Hernández 2015). In their recent redescription of *W. fieldensis* based on abundant new material housed at the Royal Ontario Museum Invertebrate Paleontology collections, Vannier et al. (2018) reported the presence of preserved CNS in two specimens (USNM 138231, ROMIP 64293; fig. 11a, b). In both cases they reported the presence of a condensed dorsal brain including direct connections with the lateral eyes via eye stalks, an esophageal foramen (the “stomodeal area” in their terminology), an anteromedian protocerebral projection of the CNS located within the anterior sclerite (the “median triangular sclerite” in their terminology), and critically, the presence of an enigmatic structure identified as the “post-protocerebral aperture” (see fig. S11a, b). Vannier et al. (2018) regarded this post-protocerebral aperture, which is located anteriorly embedded within the brain in front of the esophageal foramen and occupies a midline position relative to the stalked lateral eyes, as a legitimate feature and regarded it as evidence against the hypothesis that the euarthropod CNS is highly conserved in extant and fossil taxa (e.g. Ma et al. 2015; Strausfeld et al. 2016). In the context of our paleoneuranatomical observations in *Anacheirurus adserai* (MCZ:IP:202503; Fig. 1; fig. S1), we put forward a reinterpretation of *W. fieldensis* in which the enigmatic post-protocerebral aperture (*sensu* Vannier et al. 2018) most likely represents evidence of the central complex in this stem-group mandibulate given their fundamentally similar location on the brain and between the protocerebral (ocular) lobes. The difference in the morphology of the central complex in *W. fieldensis*, which has a flattened and elongate appearance, relative to that of *A. adserai*, which is furcula-like shaped, falls comfortably within the variability observed among extant mandibulates (e.g. Homberg 2008; Thoen et al. 2017). Given that the external exoskeletal and appendicular morphology of *W. fieldensis* supports affinities with stem-group Mandibulata, we posit that our reinterpretation for the presence of central complex in the CNS of this taxon is more parsimonious and congruent with the known neuroanatomy of extant and extinct euarthropods without the need to invoke a unique or enigmatic nature for this structure.

A similar midline structure has also been observed in *Odaraia alata* (Ortega-Hernández 2015), particularly in specimen USNM 189232 (fig. 11c, d). The head of *O. alata* features a well-developed anterior sclerite that contains a portion of the CNS that mostly encompasses the area of the brain between the lateral eye stalks and bifurcates posteriorly into the esophageal foramen. The anteriormost portion of the preserved CNS in USNM 189232 shows three highly reflective structures, two in an anterolateral position and one median located immediately posterior to the former, that have been previously identified as the so-called “frontal organs”, which most likely represent simple median eyes. Critically, median eyes/frontal organs in Cambrian euarthropods are typically paired as also observed in a another well preserved specimen of *O. alata* (see Ortega-Hernández 2015), so the presence of a third median structure in USNM 189232 deviates from this general pattern. Likewise, recent work by Schoenemann and Clarkson (2023) has argued that two median eyes correspond to the ancestral organization for crown-group euarthropods. The similarity in terms of position relative to the stalked lateral eyes and the esophageal foramen and considering the new observations from *Anacheirurus adserai* (Fig. 1; fig. S1) and *Waptia fieldensis* (fig. S11a, b), we propose that USNM 189232 also most likely shows evidence for a morphologically simple central complex in *O. alata*. Ultimately, the existence of a central complex in fossil euarthropods is to be expected given that this midline neuropil represents a fundamental component of the protocerebrum in extant representatives, albeit with a significant degree of morphological variability and complexity (e.g. Homberg et al. 2008; Loesel et al. 2013; Thoen et al. 2017). The recognition of the central complex in Cambrian and Ordovician euarthropods, however, has proved more challenging given that this neuropil is embedded within the brain and thus easily concealed during fossilization, particularly because fossil CNS are mostly recognizable based on the morphology of their outlines (e.g. Ma et al. 2012, 2015; Tanaka et al. 2013; Cong et al. 2014; Vannier et al. 2018; Ortega-Hernández et al. 2019, 2022; Bicknell et al. 2021). These comparisons and our new data open the possibility for an improved search image for fine neuroanatomical structures preserved in the fossil record that can provide additional information to reconstruct the evolution of the CNS in deep time.

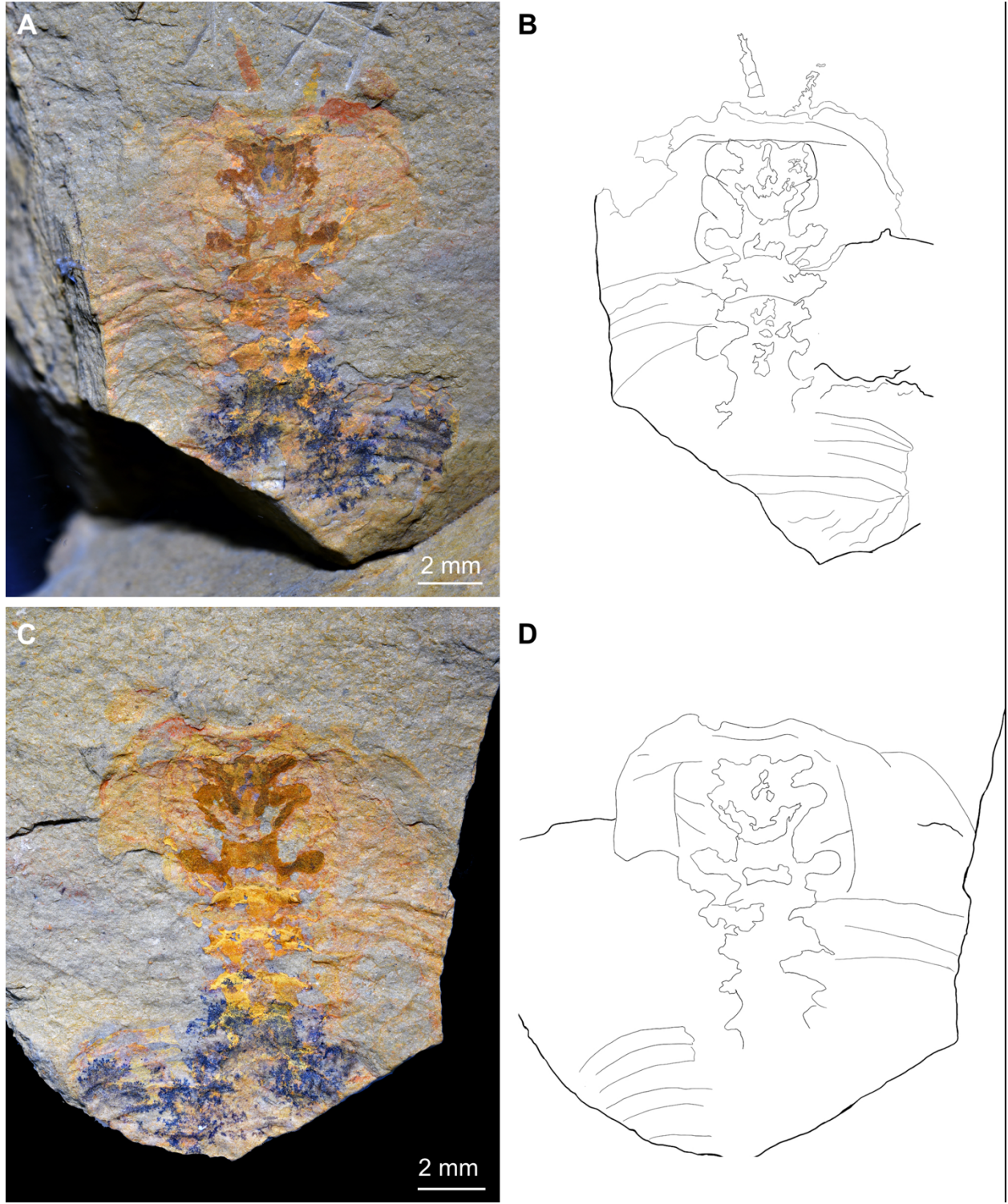

**Fig. S1. Central nervous system in the phacopid trilobite *Anacheirurus adserai* from the Lower Ordovician Fezouata Shale biota of Morocco. a, MCZ:IP:202503a, part. b,**

MCZ:IP:202503b, counterpart. Institutional abbreviations: MCZ, Museum of Comparative Zoology, USA. **c**, Low magnification of MCZ:IP:202503a, part.

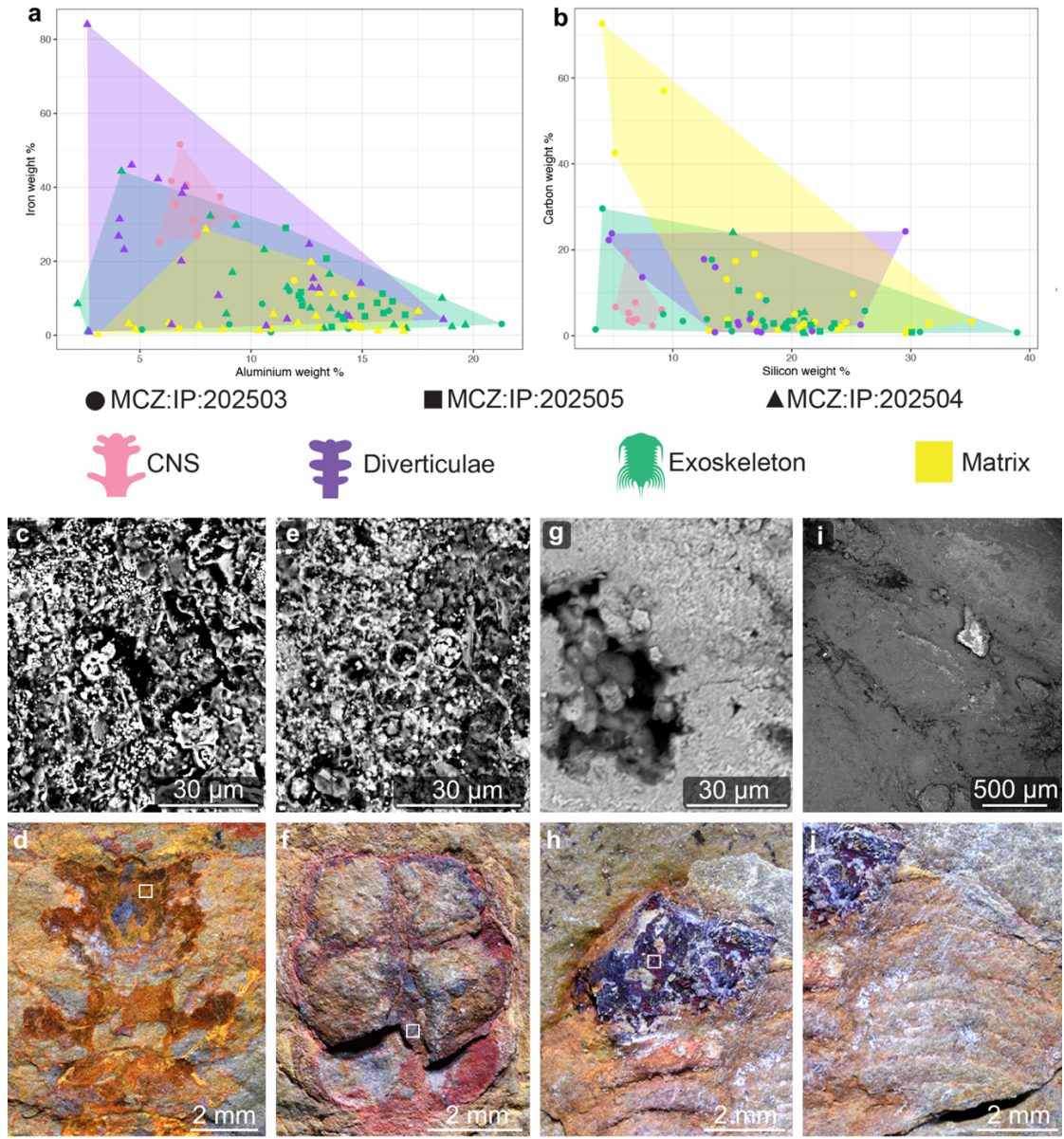

**Fig. S2. Comparative elemental analyses of nervous and digestive system preservation in *Anacheirurus* from the Fezouata Shale. **a, b**, Bivariate graphs of elemental weight percent from SEM-EDS spot analyses (see Extended Data Figs. 2-4). **c**, Backscattered electron micrograph of nervous system in MCZ:IP:202503 showing framboidal iron deposition. **d**, Nervous system of *Anacheirurus adserai* (MCZ:IP:202503) with box showing location of panel c. **e**, Backscattered electron micrograph of cephalic gut diverticulae in MCZ:IP:202504 showing framboidal iron deposition. **f**, Cephalic gut diverticulae of *Anacheirurus* sp. (MCZ:IP:202504) with box showing location of panel e. **g**, Backscattered electron micrograph showing euhedral iron overgrowth on glabella of MCZ:IP:202505. **h**, Glabella with diagenetic iron overgrowth in *Anacheirurus adserai* (MCZ:IP:202505) with box showing location of panel g. **i**, Backscattered electron micrograph**

showing pleurae of MCZ:IP:202505 without significant iron deposition. **j**, Pleurae of *Anacheirurus adserai* (MCZ:IP:202505) with box showing location of panel i.

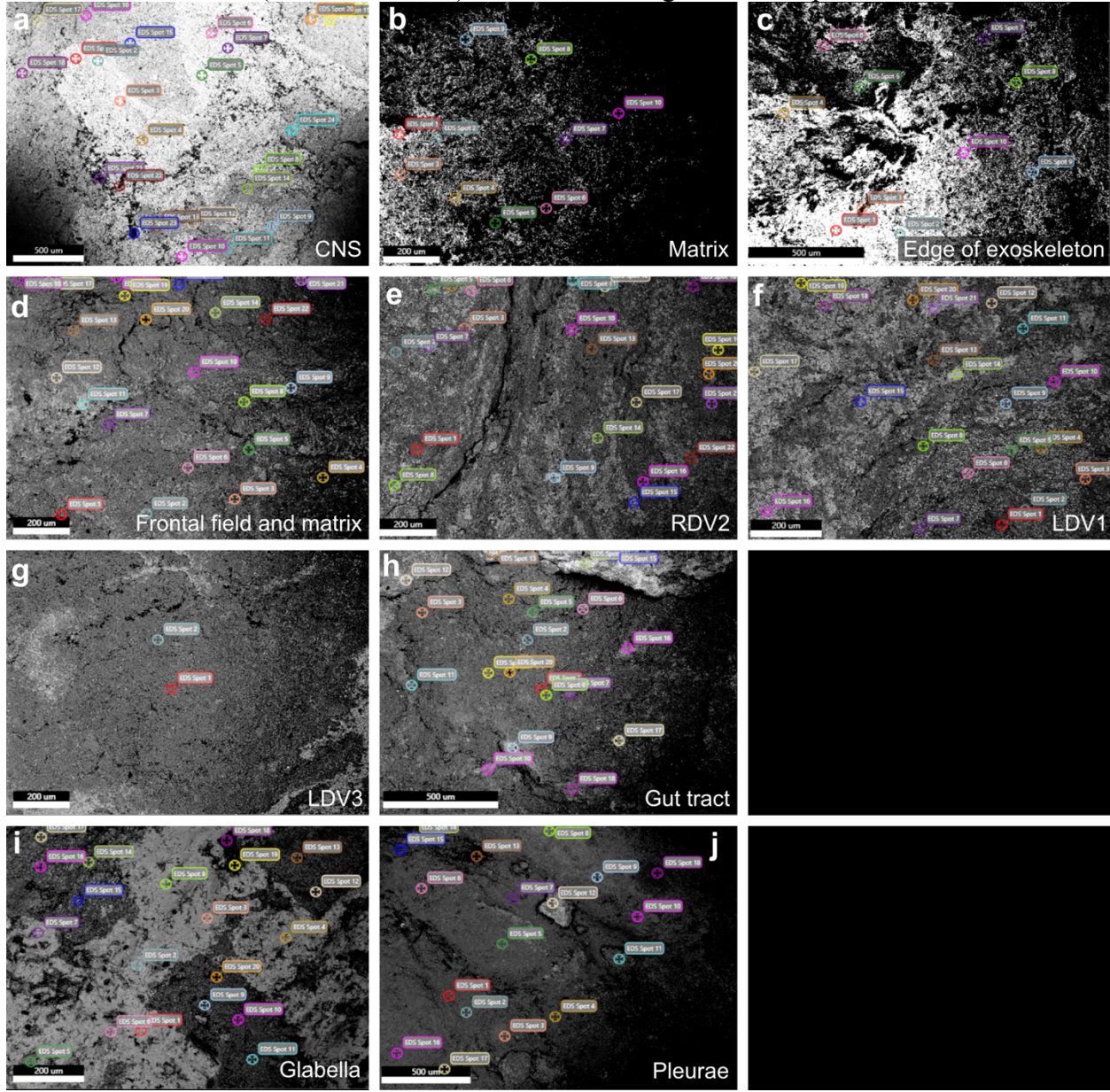

**Fig. S3. Spot analyses of *Anacheirurus* from the Fezouata Shale. a – c, *Anacheirurus adserai* (MCZ:IP:202505). a, Location of spot analyses on CNS. b, Location of spot analyses in matrix anterior of the fossil. c, Location of spot analyses at the edge of the exoskeleton. d – h, *Anacheirurus* sp. (MCZ:IP:202504). d, Location of spot analyses on frontal field and matrix. e, Location of spot analyses on right second diverticulae. f, Location of spot analyses on left first diverticula. g, Location of spot analyses on left third diverticula. h, Location of spot analyses on gut tract. i, j, *Anacheirurus adserai* (MCZ:IP:202503). i, Location of spot analyses on glabella. j, Location of spot analyses on pleurae.**

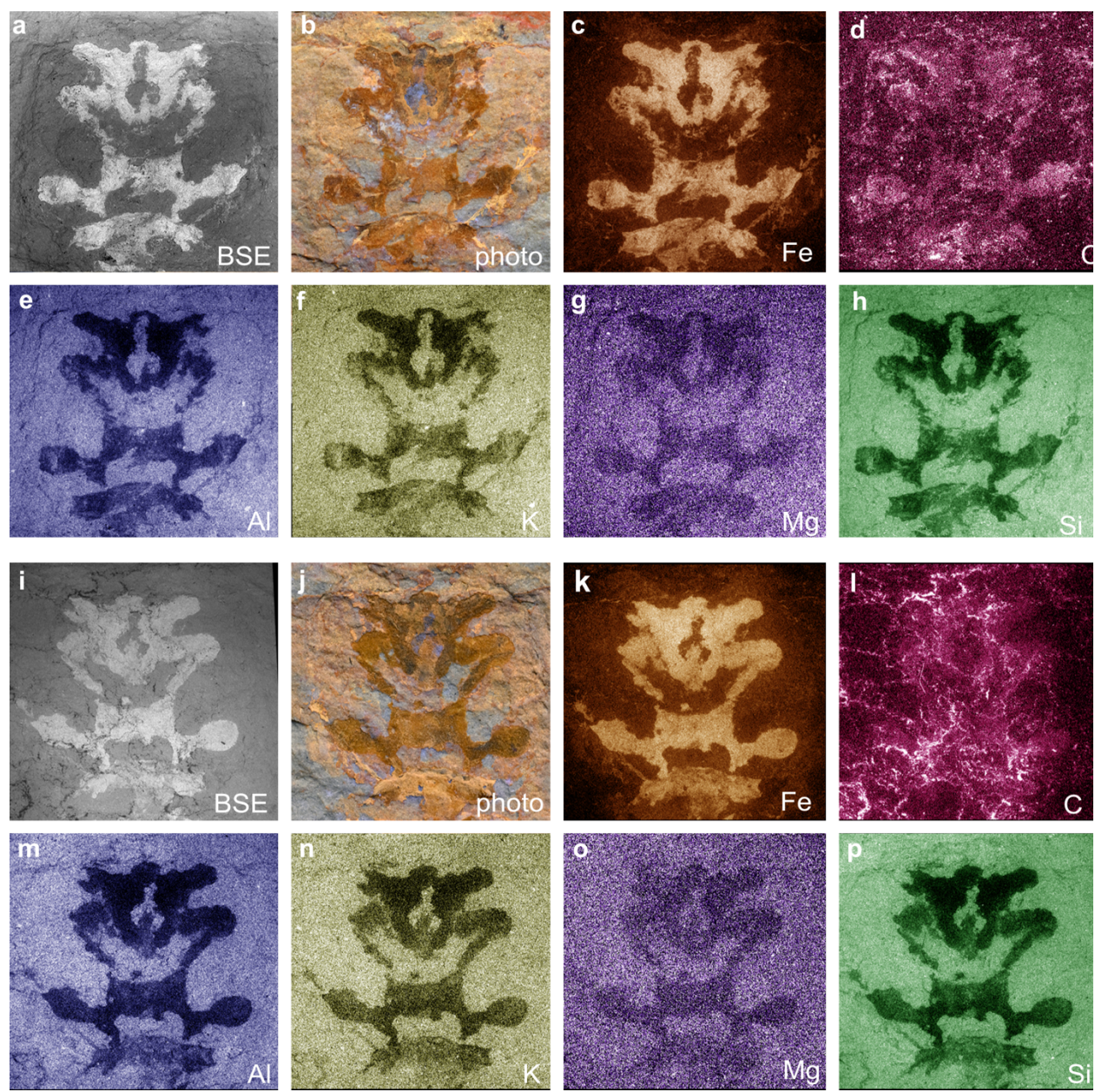

**Fig. S4. Elemental composition of the central nervous system in *Anacheirurus adserai* MCZ:IP:202523. a – g, Part. h – n, Counterpart.**

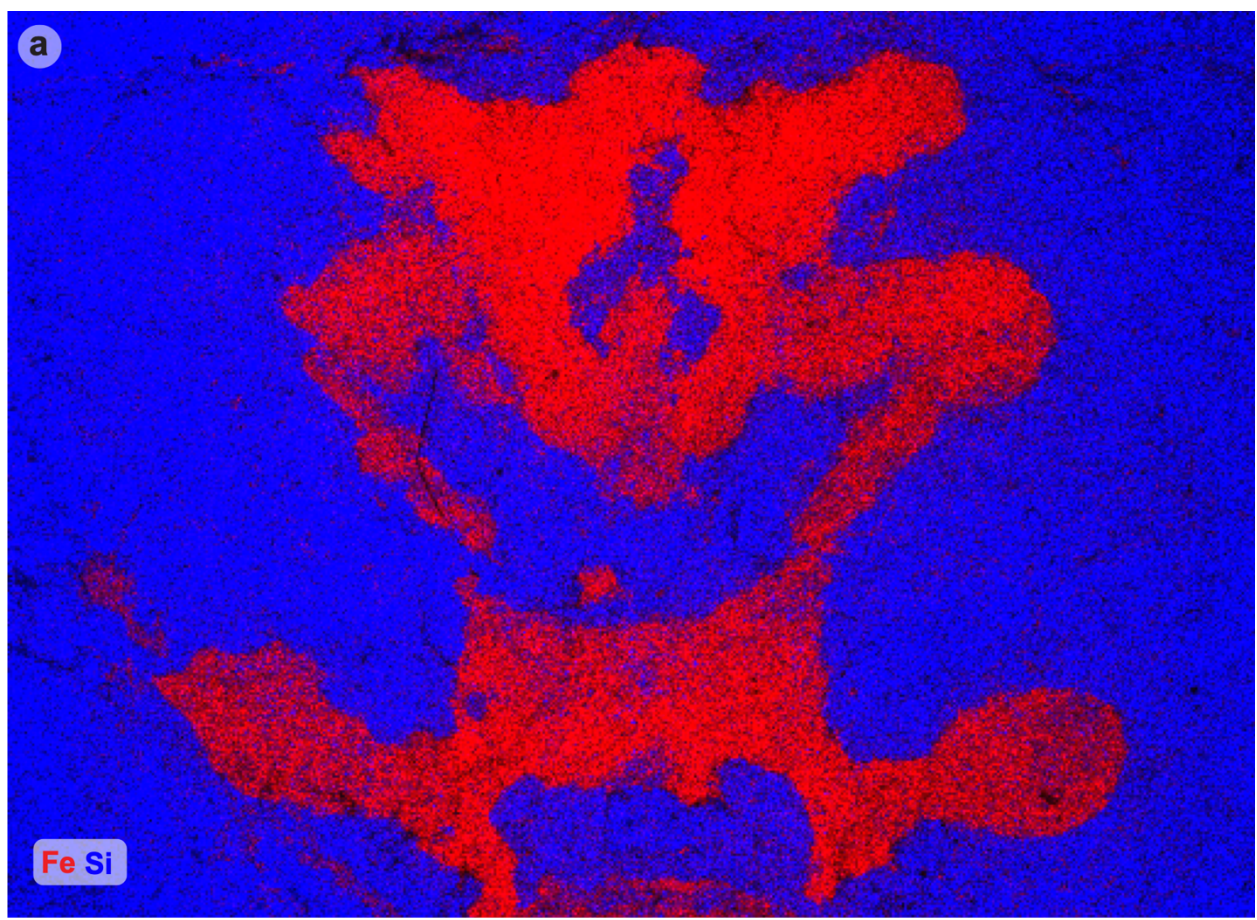

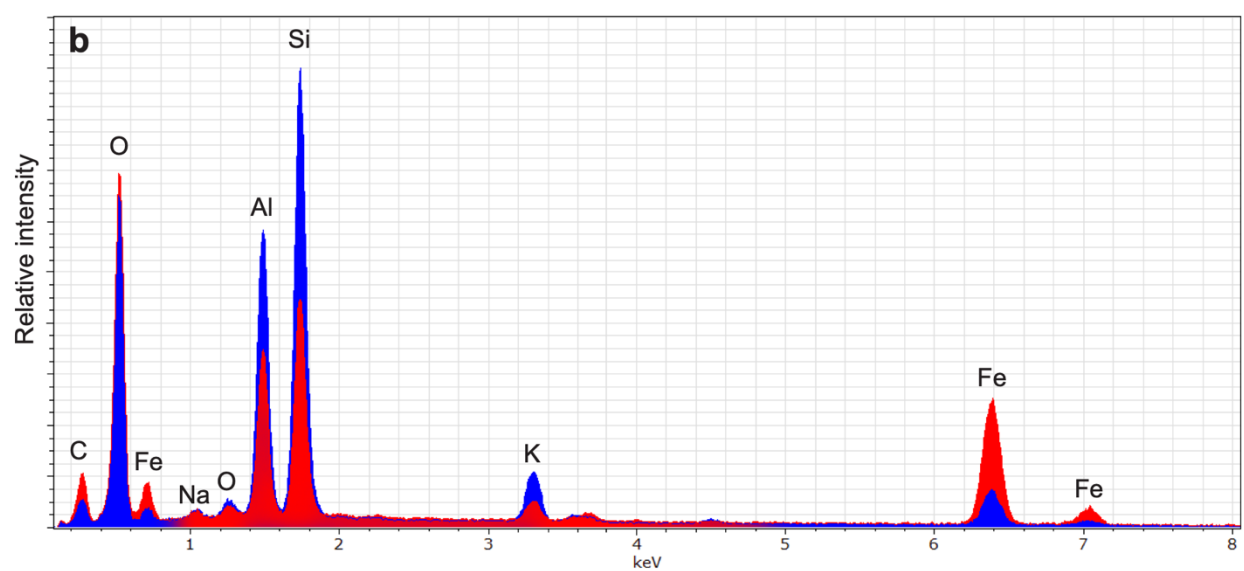

**Fig. S5. Elemental composition of *Anacheirurus adserai* MCZ:IP:202503. a, EDS silicon and iron map of counterpart. b, Representative EDS spectra of blue and red phases from a.**

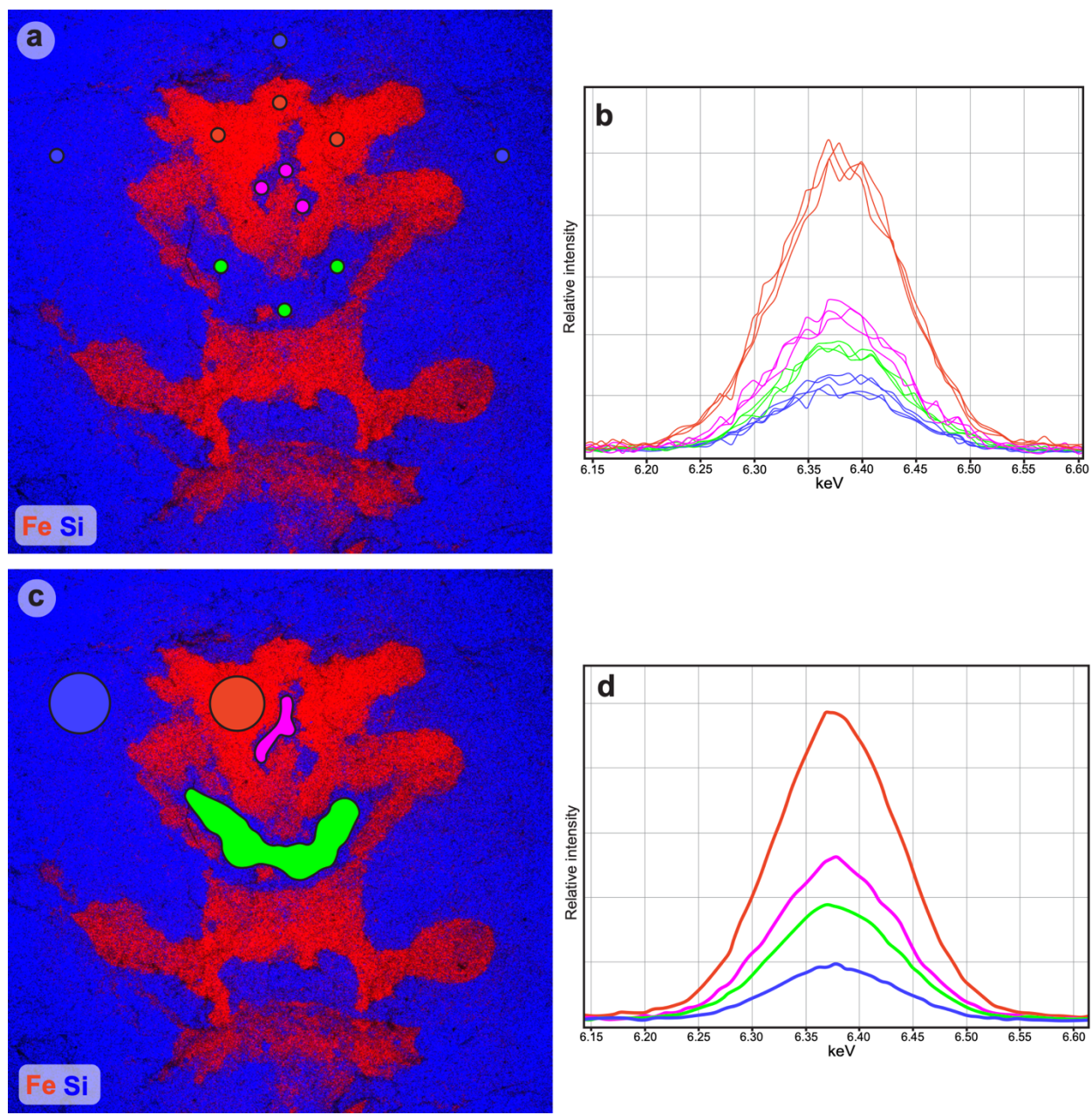

**Fig. S6. Iron concentration differences in *Anacheirurus adserai* MCZ:IP:202503 counterpart.** Red, nervous system; magenta, central complex; green, esophageal foramen; blue, exoskeleton. **a**, EDS map of counterpart with locations of spot analyses marked. **b**, Individual EDS iron peaks from locations denoted in **a**. **c**, EDS map of counterpart with marked regions of interest. **d**, averaged EDS iron peaks from regions denoted in **c**.

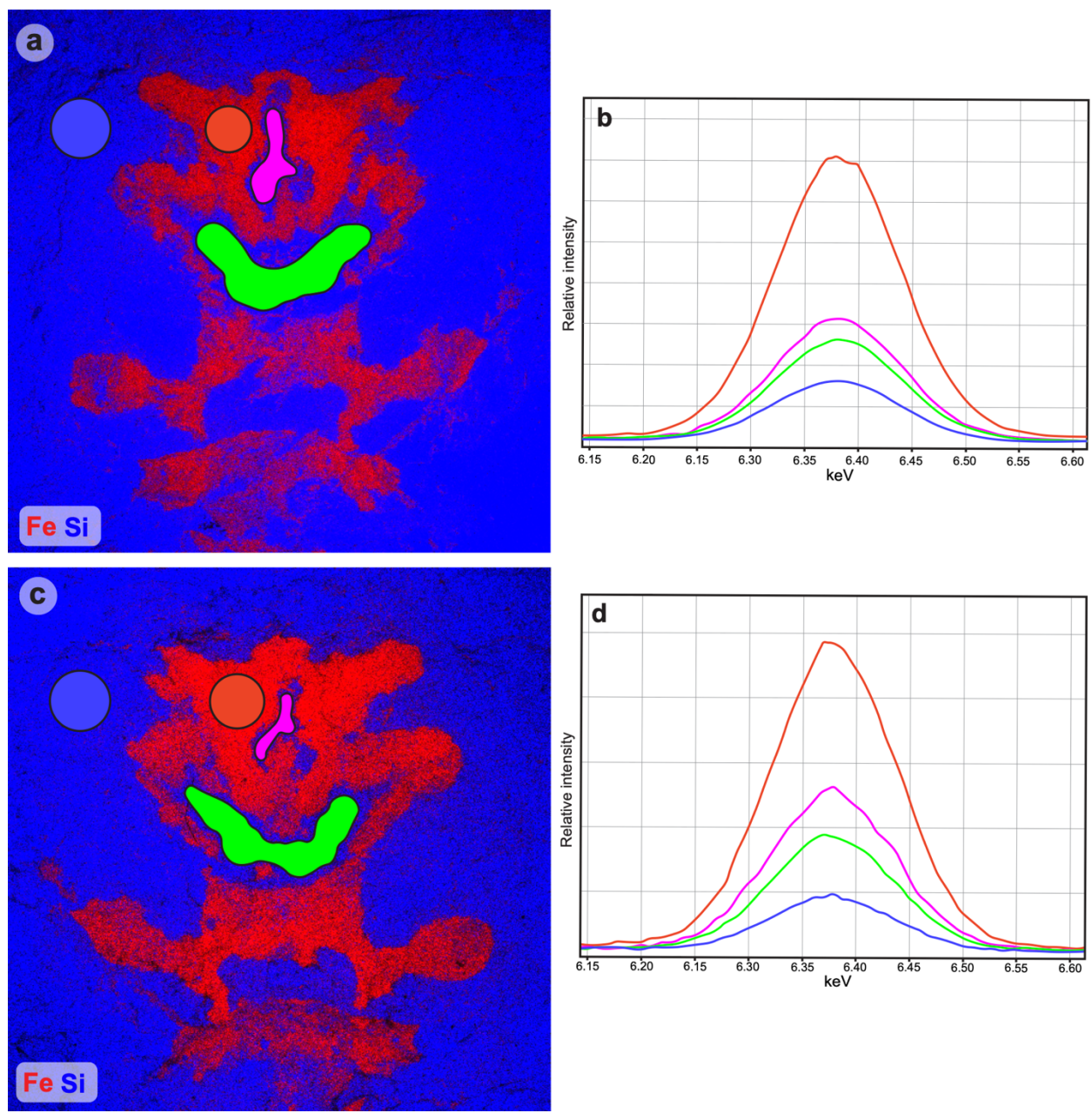

**Fig. S7. Iron concentration differences in *Anacheirurus adserai* MCZ:IP:202523 part and counterpart.** Red, nervous system; magenta, central complex; green, esophageal foramen; blue, exoskeleton. **a, b**, Part. **c, d**, counterpart.

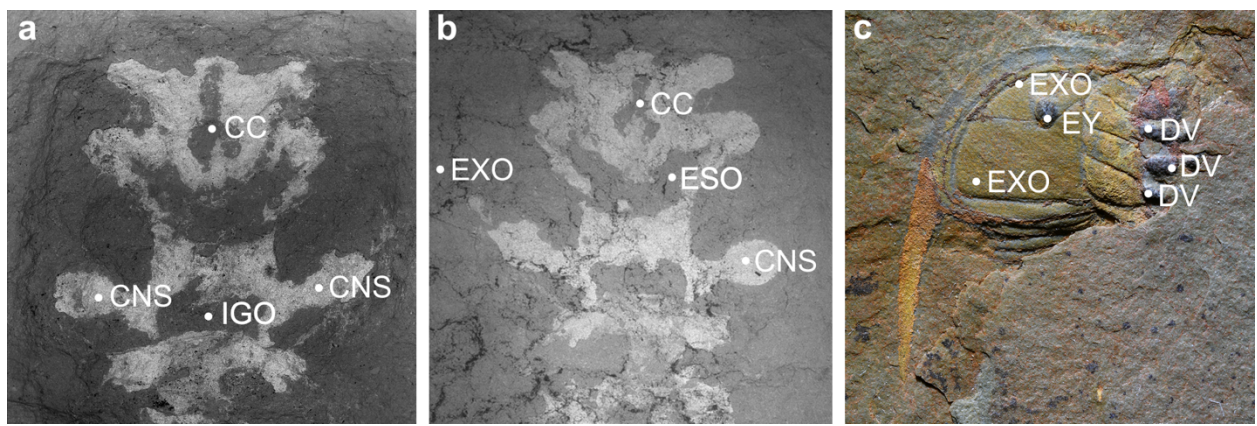

**Fig. S8. Spot analyses of *Anacheirurus* from the Fezouata Shale. a – c, *Anacheirurus adersai* (MCZ:IP:202505). a, Location of spot analyses on CNS. b, Location of spot analyses in matrix anterior of the fossil. c, Location of spot analyses at the edge of the exoskeleton. d – h, *Anacheirurus* sp. (MCZ:IP:202504). d, Location of spot analyses on frontal field and matrix. e, Location of spot analyses on right second diverticulae. f, Location of spot analyses on left first diverticula. g, Location of spot analyses on left third diverticula. h, Location of spot analyses on gut tract. i, j, *Anacheirurus adserai* (MCZ:IP:202503). i, Location of spot analyses on glabellae. j, Location of spot analyses on pleurae. Abbreviations: CC, central complex; CNS, Central nervous system; DV, diverticulae; ESO, esophageal foramen; EXO, exoskeleton; EY, eye.**

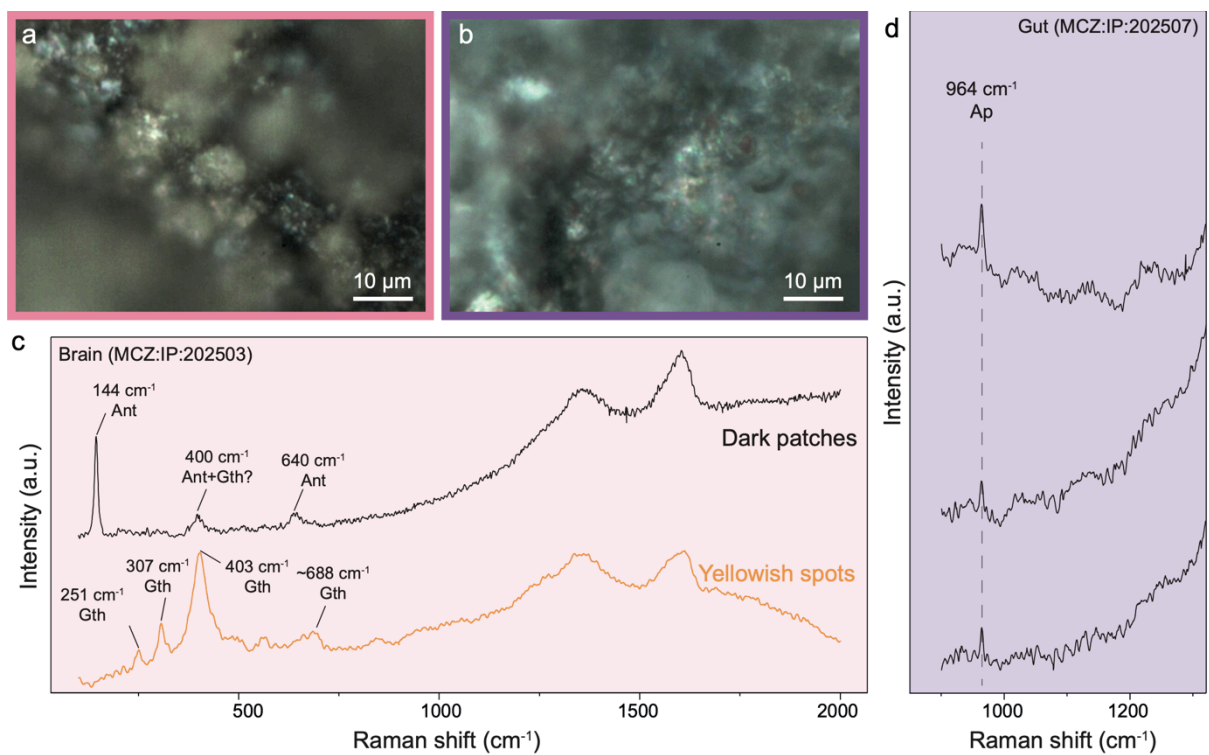

**Fig. S9. Light photographic images and Raman spectra from the CNS and gut tissues. a,** Reflected light image of the CNS region in MCZ:IP:202503. **b,** Reflected light image of the gut region in MCZ:IP:202507. **c,** Representative spectra from dark, carbon-rich regions and yellowish spots in the CNS. **d,** Zoomed-in spectra showing the 964 cm<sup>-1</sup> apatite peak in gut tissue.

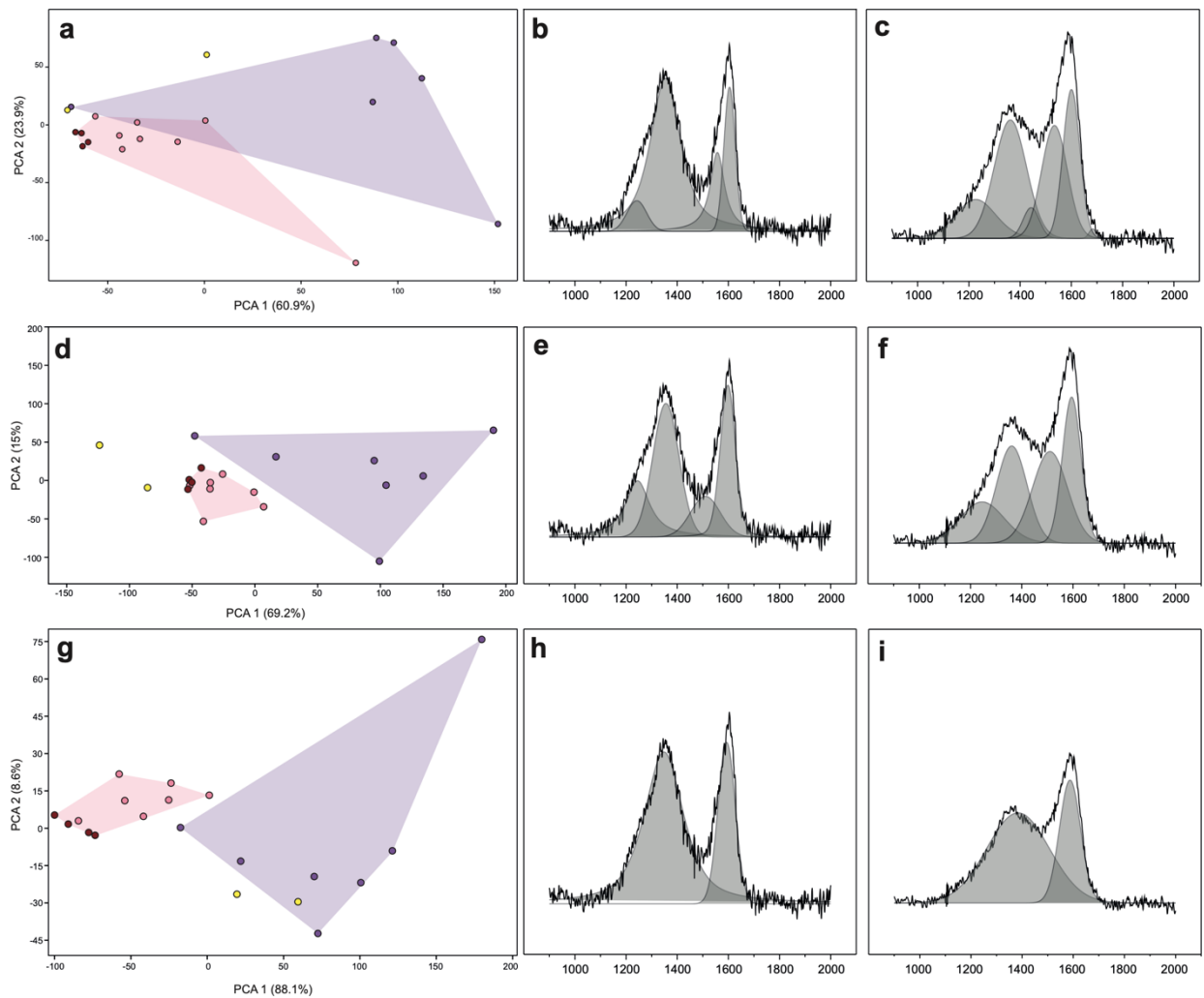

**Fig. S10. Principal Component Analysis (PCA) plots highlighting variations in the Raman band features and their respective fitting procedures.** **a, d, g,** PCA from the unconstrained fitting and representative spectra from the CNS (a2) and gut tissues (a3), with the fitted line functions. **b, e, h,** PCA plots from the ‘Fitting G’ protocol of Kouketsu et al. (2014) and respective fitted line functions. **c, f, i,** PCA plots from the minimal-component fitting and its respective line functions in the representative spectra.

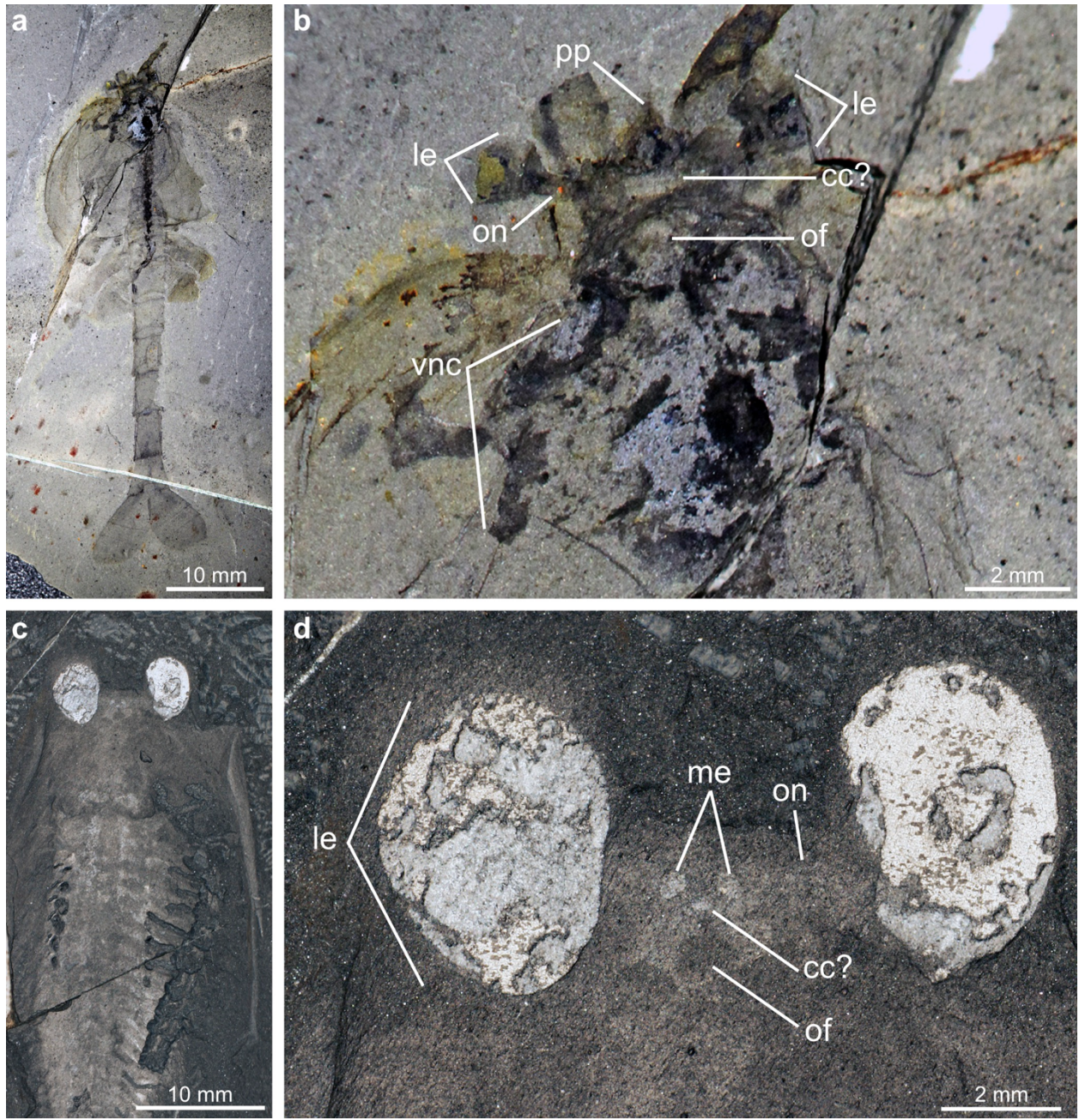

**Fig. S11. Central nervous system preservation in stem-group mandibulates from the mid-Cambrian Burgess Shale.** **a**, ROMIP 64293, *Waptia fieldensis* (courtesy of Jean-Bernard Caron). **b**, Magnification of head region in ROMIP 64293 showing fossilized brain neuroanatomy. **c**, USNM 189232, *Odaraia alata*. **d**, Magnification of head region in USNM 189232 showing fossilized brain neuroanatomy. Abbreviations: cc?, central complex; le, lateral eye; me, median eye; of, oesophageal foramen; on, optic nerve; pp, protocerebral projection, vnc, ventral nerve cord. Additional institutional abbreviations: ROMIP, Royal Ontario Museum Invertebrate Paleontology, Canada; USNM, Smithsonian National Museum of Natural History.

preparation and operating procedure. *International Journal of Coal Geology*, 191, pp. 135-151.

- Homberg, U., 2008. Evolution of the central complex in the arthropod brain with respect to the visual system. *Arthropod structure & development*, 37(5), pp.347-362.
- Kouketsu, Y., Mizukami, T., Mori, H., Endo, S., Aoya, M., Hara, H., Nakamura, D., and Wallis, S., 2014. A new approach to develop the Raman carbonaceous material geothermometer for low-grade metamorphism using peak width. *Island Arc*, 23(1), pp. 33-50.
- Loesel, R., Wolf, H., Kenning, M., Harzsch, S. and Sombke, A., 2013. Architectural principles and evolution of the arthropod central nervous system. *Arthropod biology and evolution: molecules, development, morphology*, pp.299-342.
- Ma, X., Hou, X., Edgecombe, G.D. and Strausfeld, N.J., 2012. Complex brain and optic lobes in an early Cambrian arthropod. *Nature*, 490(7419), pp.258-261.
- Ma, X., Edgecombe, G.D., Hou, X., Goral, T. and Strausfeld, N.J., 2015. Preservation pathways of corresponding brains of a Cambrian euarthropod. *Current Biology*, 25(22), pp.2969-2975.
- Murad, E., 1997. Identification of minor amounts of anatase in kaolins by Raman spectroscopy. *American Mineralogist*, 82(1-2), pp. 203-206.
- Ortega-Hernández, J., 2015. Homology of head sclerites in Burgess Shale euarthropods. *Current Biology*, 25(12), pp.1625-1631.
- Ortega-Hernández, J., Leroosey-Aubril, R., Losso, S.R. and Weaver, J.C., 2022. Neuroanatomy in a middle Cambrian mollisoniid and the ancestral nervous system organization of chelicerates. *Nature communications*, 13(1), p.410.
- Potin, G. J.-M., Gueriau, P. and Daley, A.C., 2023. Radiodont frontal appendages from the Fezouata Biota (Morocco) reveal high diversity and ecological adaptations to suspension-feeding during the Early Ordovician. *Frontiers in Ecology and Evolution*, 11, 1214109.
- Rossi, V., Unitt, R., and McNamara, M., 2024. A new non-destructive method to decipher the origin of organic matter in fossils using Raman spectroscopy. *RSC advances*, 14(37), 26747-26759.
- Saleh, F. et al., 2020a. Taphonomic pathway of exceptionally preserved fossils in the Lower Ordovician of Morocco. *Geobios*, 60, pp. 99–115.

- Schoenemann, B. and Clarkson, E.N., 2023. The median eyes of trilobites. *Scientific Reports*, 13(1), p.3917.
- Strausfeld, N.J., Ma, X., Edgecombe, G.D., Fortey, R.A., Land, M.F., Liu, Y., Cong, P. and Hou, X., 2016. Arthropod eyes: the early Cambrian fossil record and divergent evolution of visual systems. *Arthropod Structure & Development*, 45(2), pp.152-172.
- Tanaka, G., Hou, X., Ma, X., Edgecombe, G.D. and Strausfeld, N.J., 2013. Chelicerate neural ground pattern in a Cambrian great appendage arthropod. *Nature*, 502(7471), pp.364-367.
- Thoen, H.H., Marshall, J., Wolff, G.H. and Strausfeld, N.J., 2017. Insect-like organization of the stomatopod central complex: functional and phylogenetic implications. *Frontiers in Behavioral Neuroscience*, 11, p.12.
- Vannier, J., Aria, C., Taylor, R.S. and Caron, J.B., 2018. *Waptia fieldensis* Walcott, a mandibulate arthropod from the middle Cambrian Burgess Shale. *Royal Society open science*, 5(6), p.172206.
